# Quantification of archaea-driven freshwater nitrification: from single cell to ecosystem level

**DOI:** 10.1101/2021.07.22.453385

**Authors:** Franziska Klotz, Katharina Kitzinger, David Kamanda Ngugi, Petra Büsing, Sten Littmann, Marcel M. M. Kuypers, Bernhard Schink, Michael Pester

## Abstract

Deep oligotrophic lakes sustain large archaeal populations of the class *Nitrososphaeria* in their hypolimnion. They are thought to be the key ammonia oxidizers in these freshwater systems and as such responsible for the rate-limiting step in nitrification. However, the impact that planktonic *Nitrososphaeria* have on N cycling in lakes is severely understudied and yet to be quantified. Here, we followed this archaeal population in one of Central Europe’s largest lakes, Lake Constance, over two consecutive years using metagenomics and metatranscriptomics combined with stable isotope-based activity measurements. A single, highly abundant and transcriptionally active freshwater ecotype of *Nitrososphaeria* dominated the nitrifying community. Phylogenomic analysis of its metagenome-assembled genome showed that this ecotype represents a new lacustrine *Nitrosopumilus* species. Stable isotope probing revealed that *Nitrososphaeria* incorporated significantly more ^15^N-labeled ammonium than most other microorganisms at near-natural conditions and oxidized ammonia at an average rate of 0.22 ± 0.11 fmol cell^−1^ d^−1^. This translates to 1.9 gigagram of ammonia oxidized per year, corresponding to 12% of the N-biomass produced annually by photosynthetic organisms in Lake Constance. Here, we show that ammonia-oxidizing archaea play an equally important role in the nitrogen cycle of deep oligotrophic lakes as their counterparts in marine ecosystems.

## Introduction

Freshwater lakes are important drinking water reservoirs. To be suitable for drinking water and to prevent toxicity for fish, ammonia must not accumulate. Nitrification prevents an accumulation of ammonia and converts it to nitrate via nitrite, with ammonia oxidation being the rate-limiting step^1^. Although nitrification does not directly change the inventory of inorganic N in freshwater ecosystems, it represents a critical link between mineralization of organic N and its eventual loss as N_2_ to the atmosphere by denitrification or anaerobic ammonium oxidation^1^. The process of ammonia oxidation is catalyzed by three different microbial guilds. Two of these, the ammonia-oxidizing archaea (AOA)^2,3^ and the ammonia-oxidizing bacteria (AOB)^4^ oxidize ammonia to nitrite and depend on nitrite-oxidizing bacteria (NOB)^5^ to complete nitrification by further oxidation of nitrite to nitrate. The third guild oxidizes ammonia directly to nitrate and is therefore referred to as complete ammonia oxidizers (comammox)^6,7^.

In general, the ratio of AOA to AOB decreases with increasing trophic state of freshwater lakes, as AOB have a preference for increased inorganic nitrogen loading and AOA are sensitive towards copper complexation by organic matter^8–14^. In contrast, comammox bacteria were detected at very low abundances in lacustrine systems, if at all^15,16^. In snapshot analyses of oligotrophic lakes, AOA typically outnumbered AOB, especially in the deep oxygenated hypolimnion, and constituted up to 19% of the archaeal and bacterial picoplankton^10,15,17–20^. These observations resemble the situation in marine ecosystems, where AOA account for up to 40% of all microorganisms in the deep sea and thus are estimated to be among the most numerous microorganisms on earth^21,22^. In contrast to marine ecosystems, the impact that planktonic freshwater AOA have on N cycling in oligotrophic lakes, which are often important drinking water reservoirs, is severely understudied and yet to be quantified.

Lake Constance is an oligotrophic, fully oxygenated lake that provides drinking water to more than five million people^23^. Being the second largest lake by volume in Central Europe, it represents an important model habitat for limnological processes^24^. A year-round survey of Lake Constance waters established that a single 16S rRNA gene-ecotype of AOA, related to the genus *Nitrosopumilus* (class *Nitrososphaeria*, formerly known also as Thaumarchaeota^25^), constituted the largest nitrifier population along the depth profile throughout the year, with AOB being two orders of magnitude less abundant^16^. At the same time, comammox bacteria were below the detection limit as based on diagnostic PCR of the *amoA* gene^16^, which codes for the structural subunit of the key enzyme for ammonia oxidation, ammonia monooxygenase. Here, we link the large prevailing AOA population in this oligotrophic lake with nitrification activity and quantify this important ecosystem service at the single-cell, population, and ecosystem level.

## Results & Discussion

### AOA account for 8-39% of total archaea and bacteria in the hypolimnion

Depth profiles of total ammonium (NH_4_^+^ + NH_3_) and nitrate showed typical inverse concentration profiles during periods of primary productivity (Fig. 1a-c). Total ammonium decreased and nitrate increased towards the hypolimnion, indicative of active ammonium consumption and nitrate production in hypolimnetic waters, presumably by nitrification (Fig. 1b,c). Hence, we decided to follow annual dynamics of the AOA population in the central part of the hypolimnion at 85 m depth. Copy numbers of Lake Constance-specific archaeal *amoA* genes were enumerated by quantitative PCR (qPCR). As *amoA* is typically present as a single copy gene in AOA^26^, *amoA* copy numbers were compared to total archaeal and bacterial 16S rRNA gene copy numbers to estimate AOA relative abundance. This confirmed the presence of a large prevailing AOA population in the hypolimnion ranging from 8.0 ± 0.9% to 38.9 ± 5.7% of the archaeal and bacterial picoplankton. Absolute abundances ranged between 9.8 × 10^3^ ± 1.0 × 10^4^ and 1.2 × 10^5^ ± 4.9 × 10^4^ *amoA* copies ml^−1^ in winter and summer, respectively, with a mean copy number of 4.3 × 10^4^ *amoA* copies ml^−1^ (Fig. 1d). These qPCR results corresponded well to direct *Nitrososphaeria* cell counts based on catalyzed reporter deposition fluorescence *in situ* hybridization (CARD-FISH) at two selected time points (Supplementary Fig. 1). We therefore use the terms AOA and *Nitrososphaeria* as synonyms from here on.

**Figure 1.**
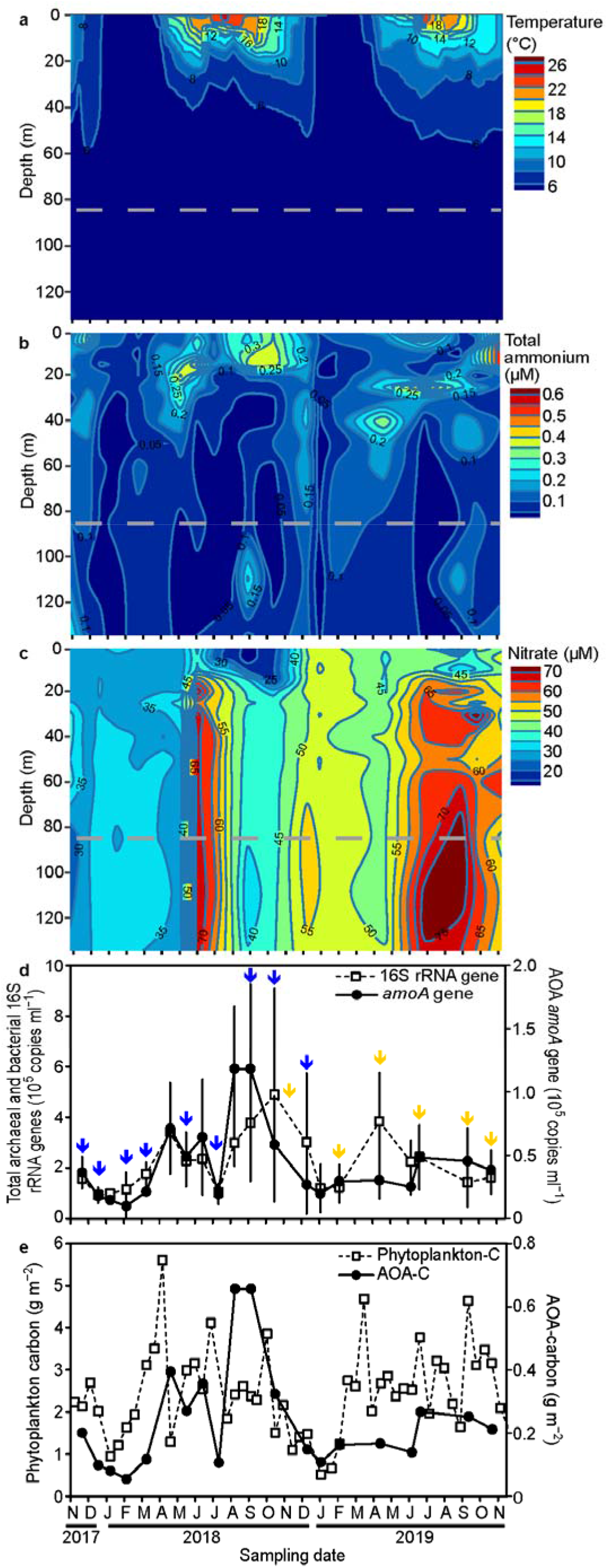
Seasonality of physico-chemical parameters (a-c), the total archaeal and bacterial picoplankton as well as the AOA population at 85 m depth (d), and the carbon budget of the phytoplankton and AOA population (e) over two consecutive years in the water column of Lake Constance. The sampling depth for qPCR, metagenome and metatranscriptome analyses is indicated as a dashed grey line at 85 m. Arrows indicate time points for metagenome (blue) and metatranscriptome (yellow) sampling. qPCR analyses were done in replicates of 3 or 4, except for December 18^th^, 2018, where only duplicates could be measured for the AOA *amoA*.

### A freshwater-specific AOA species dominates the nitrifier community

To gain insight into the genetic repertoire of the AOA, we conducted a metagenomic survey of hypolimnetic waters covering nine time points from Nov 2017 to Dec 2018 (Fig. 1d). Best assembly results were obtained by a co-assembly of the winter datasets Nov 2017, Dec 2017 and Feb 2018, which resulted in a single metagenome-assembled genome (MAG) related to *Nitrososphaeria* that was further refined by long PacBio reads for scaffolding. This resulted in a high quality assembly of eight contigs spanning 1.2 Mbp, with a checkM-estimated^27^ coverage of 99% and contamination of 0% (Supplementary Table 1). The 16S rRNA gene of MAG AOA-LC4 was 100% identical to the single 16S rRNA gene-ecotype that was previously shown in an amplicon-based study to outnumber all other AOA and AOB populations by at least two orders of magnitude throughout the depth profile of Lake Constance^16^ and thus represents the main ammonia oxidizer of this lake. An index of replication (iREP)^28^ analysis indicated active replication of the AOA-LC4 population throughout the entire year with average values of 50 ± 19% and ranging from a minimum of 34% in November/December 2017 to a maximum of 90% in October 2018 (Supplementary Table 1). Although the iRep algorithm was recently questioned to work with population genomes integrating over many co-occurring but heterogeneous close relatives^29^, it performs well for uniform population genomes such as pure cultures^28,30,31^. We consider the extremely low and highly skewed AOA diversity in Lake Constance^16^ to be rather representative of the latter case. Phylogenomic analysis placed AOA-LC4 within a freshwater/brackish water-associated clade of the *Nitrosopumilaceae*, which was distinct from marine representatives (Supplementary Fig. 2). Closest relatives were MAGs retrieved from Lake Baikal (G182) and from the Caspian Sea (Casp-thauma1), which, together with AOA-LC4, represent a new species within the genus *Nitrosopumilus* based on genome-wide average nucleotide and amino acid identities (Supplementary Fig. 3). Therefore, AOA-LC4 is a representative of AOA in major inland water bodies.

AOA-LC4 encoded the core genetic repertoire of *Nitrosopumilus* species^32,33^. This included all genes necessary for oxidation of ammonia to hydroxylamine in the canonical arrangement *amoBCxA*, as well as genes with a proposed function in further oxidation of hydroxylamine to nitrite, i.e. genes encoding (putative) multicopper oxidases including a postulated reversely operating nitrite reductase (*“nirK”*)^34,35^. In addition, AOA-LC4 carries genes for a urea transporter (*dur3*), urea amidohydrolase (*ureABC*), and urease accessory proteins (*ureDEFGH*). Furthermore, it is prototrophic for the vitamins Thiamine (B1), Riboflavin (B2), Biotin (B7) and Cobalamin (B12). For carbon fixation, it encodes the energy-efficient variant of the 3-hydroxypropionyl/4-hydroxybutyryl pathway (Supplementary Fig. 4, Supplementary Table 2).

To obtain a complete picture of the nitrifying community, we screened all obtained MAGs and single contigs for the presence of *amoA* as indicator for all ammonia oxidizers, *nxrB* (encoding the beta subunit of nitrite oxidoreductase) as indicator for all nitrite−oxidizing bacteria^36^, or both genes in the same MAG/contig as indicator for comammox bacteria^5^. MAG AOB-LC263 encoded the bacterial *amoAB* genes and was affiliated with a novel genus within the *Nitrosomonadaceae* (Proteobacteria) (Supplementary Text and Supplementary Fig. 5, 6). In addition, we found two contigs (AOB-LC199628 and AOB-LC368213), which encoded either *amoAB* or *amoCAB* and were again related to the *Nitrosomonadaceae* (Supplementary Fig. 7). However, both contigs did not group into any of the binned MAGs that fulfilled our completeness criteria (>50%). Two MAGs, NOB-LC32 and NOB-LC29, encoded *nxrAB* and *nxrB*, respectively, and were related to *Nitrospira* lineage II (Nitrospirae) (Supplementary Text and Supplementary Fig. 8‒10). Interestingly, we recovered a third *Nitrospira* lineage II MAG, COM-LC224 (Supplementary Fig. 8), which encoded both *amoA* and *nxrB* (Supplementary Fig. 10, 11). Closer inspection identified the full set of *amoCAB, hao*, and *nxrAB* encoding the ammonia monooxygenase, hydroxylamine reductase, and nitrite oxidoreductase, respectively. This finding was further supported by the affiliation of MAG COM-LC224 to comammox clade B as inferred by independent phylogenetic analysis of its concatenated single copy marker genes as well as its functional marker genes *amoA* and *nxrB* (Supplementary Text and Supplementary Fig. 8, 10, 11). Throughout the year, MAGs AOB-LC263 and COM-LC224 as well as the contigs AOB-LC199628 and AOB-LC368213 were 1–2 orders of magnitude less abundant than AOA-LC4 as inferred from mapping coverage results in the nine metagenomes (Supplementary Table 1). Similarly, MAGs NOB-LC32 and NOB-LC29 were one order of magnitude less abundant (Supplementary Table 1). This is consistent with results obtained three years earlier by 16S rRNA gene amplicon sequencing and *amoA* screening^16^.

### AOA-LC4 is transcriptionally active throughout the year

The dominance of AOA-LC4 was also reflected at the transcriptional level. In a metatranscriptome survey covering six time points from November 2018 to November 2019, *amoA* transcription of the AOA-LC4 population was consistently at least two orders of magnitude higher than of the individual AOB-LC263, AOB-LC199628, AOB-LC368213 or COM-LC224 populations (Fig. 2). Compared to *nxrB* transcription of the NOB-LC32 and NOB-LC29 populations, *amoA* transcription of AOA-LC4 was consistently one order of magnitude higher, which reflects the relative abundance estimates based on normalized metagenomic coverage (Supplementary Table 1) and 16S rRNA gene amplicons^16^. Interestingly, the transcription of the COM-LC224 *amoA* gene was one order of magnitude lower than of its *nxrB*, suggesting the possible predominance of nitrite oxidation relative to complete ammonia oxidation to nitrate in this microorganism. Alternatively, the represented two enzymes have different turnover times or regulation mechanisms.

**Figure 2.**
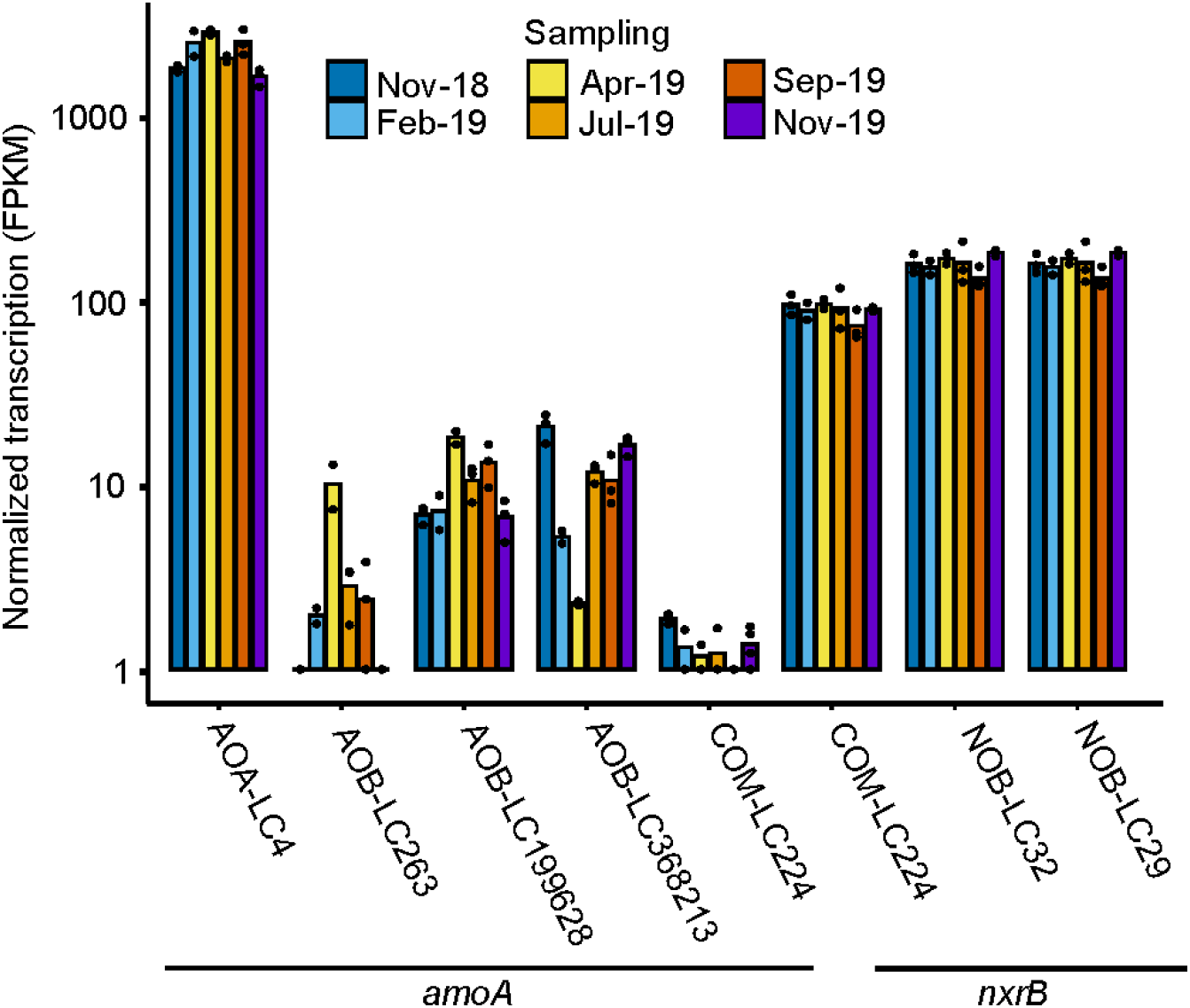
Year-round transcriptional activity of the nitrifying community in the hypolimnion (85 m depth) of Lake Constance based on metatranscriptomics. Transcription of *amoA* (encoding ammonia monooxygenase subunit A) as functional marker for all ammonia oxidizers, *nxrB* (encoding nitrite oxidoreductase subunit B) as functional marker for all nitrite oxidizers, or both genes in the same MAG as indicator for comammox bacteria is shown across all identified nitrifying microorganisms. Bars represent the mean of three replicates, except for the February and April samples, where only two replicates could by analyzed. Individual replicates are indicated by single dots. FPKM values below 1 were set to 1 for a better representation on a log_10_-scale. FPKM, fragments per kilobase of transcripts per million mapped reads.

Detailed analysis of the seasonally resolved AOA-LC4 population transcriptome identified *amoABC* among the top ten transcribed genes at a steady level with little variation throughout the yearly cycle (Fig. 3, Supplementary Table 2). Transcription of *amoABC* was highly correlated (*r_S_* ≥ 0.8) to each other and to transcription of “*nirK”* and a gene encoding a membrane-anchored PEFG-CTERM domain-containing multicopper oxidase (locus JW390_30004) (Supplementary Fig. 12). The latter two were previously postulated to be involved in the yet unresolved archaeal pathway of ammonia oxidation to nitrite^33–35,37^. On average, *amoABC* showed a 16 to 119 fold higher transcription than the highest transcribed genes encoding either proteins of the large (*rpl12*, *rpl21e*) or small ribosomal subunit (*rps11*, *rps15*), which we used as reference housekeeping genes (Fig. 3). Also *amt* and *cdvB*, encoding a high-affinity ammonium transporter and the putative cell division protein B2, respectively, were consistently found among the top ten highest transcribed genes. Since the active center of the archaeal ammonia monooxygenase is postulated to face the outside of the cytoplasmic membrane^35^, the high transcriptional levels of *amt* imply that ammonia was transported into the cell for ammonium assimilation, most likely for further biomass production as evidenced by the high transcriptional levels of *cdvB*. This is in line with the consistent transcription of genes encoding the 3-hydroxypropionate/4-hydroxybutyrate pathway for CO_2_ fixation, however, at 1 to 300 fold lower transcriptional levels as compared to the selected ribosomal reference genes (Fig. 3, Supplementary Table 2). Together, this resembles (meta-)transcriptomic data of actively growing AOA species in culture and in open marine waters with AOA-driven nitrification activity^34^. Therefore, we conclude that the AOA-LC4 population was actively oxidizing ammonia throughout the year. In contrast, genes associated with urea utilization were much less transcribed. While *dur3* (encoding an urea transporter) was transcribed similar to the four highest transcribed ribosomal protein genes, *ureABC* (encoding urease) showed a 12 to 988 fold lower transcription. The same was true for *ureDEFGH* (encoding [putative] urease accessory proteins) with transcriptional levels either below the detection limit, or 43 to 1,507 fold lower. However, low transcription of urease and urease-associated genes has previously been shown to be a poor predictor for lack of urea-based nitrification of AOA^34^.

**Figure 3:**
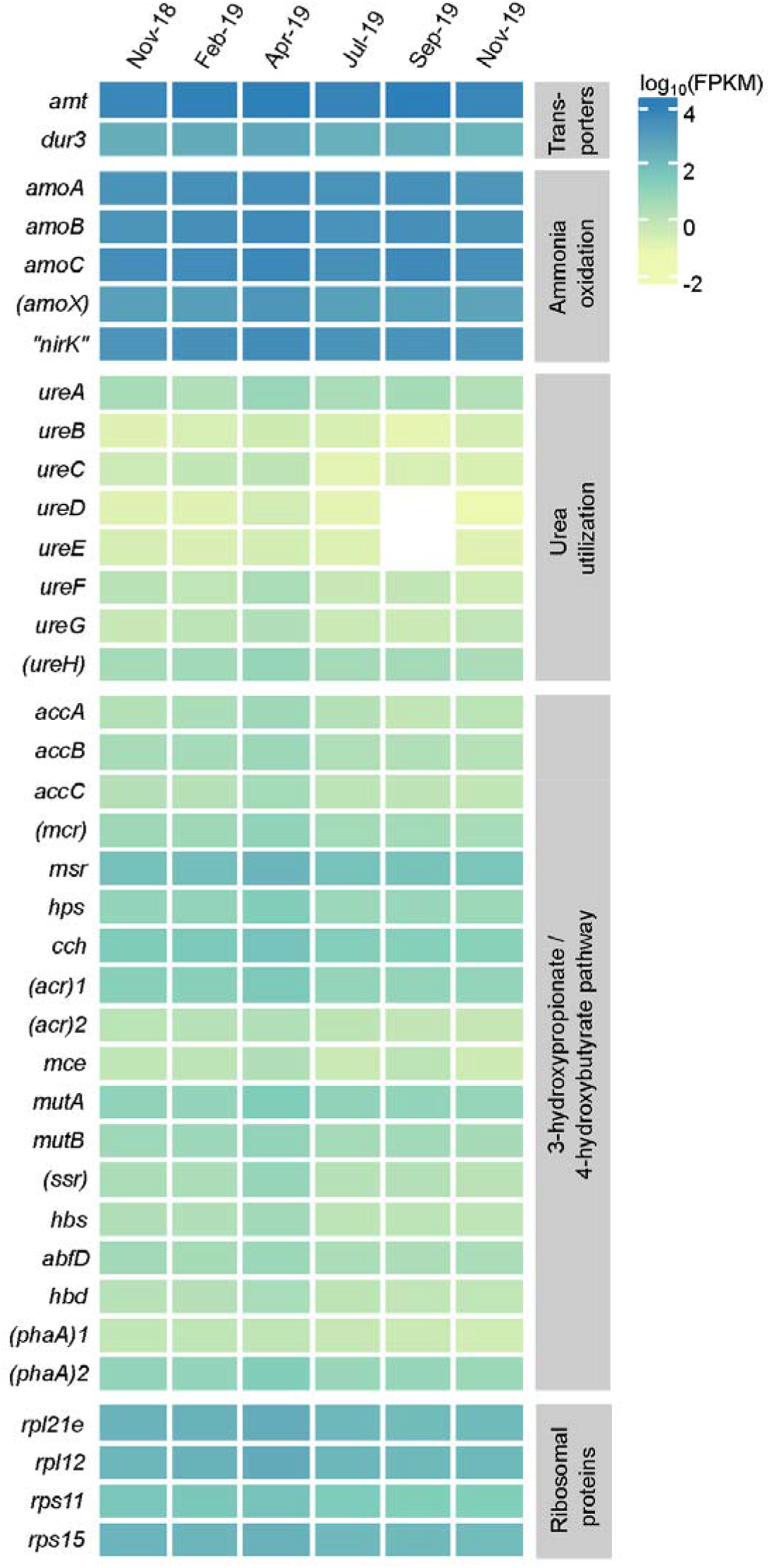
Seasonally-resolved transcription of genes involved in the nitrogen and carbon metabolism of AOA-LC4 at 85 m depth. As reference housekeeping genes, the two highest transcribed genes encoding either proteins of the large (*rpl12, rpl21e*) or small ribosomal subunit (*rps11*, *rps15*) are shown at the bottom. Transcription values are represented as log_10_-transformed mean FPKM (fragments per kilobase of transcripts per million mapped reads) values for every time point. Values represent the mean of three replicates, except for the February and April samples, where only two replicates could by analyzed. No transcription of a specific gene is indicated in white. Putatively annotated genes are given in brackets; gene duplicates were consecutively numbered.

### AOA predominate ^15^N-ammonium assimilation in the hypolimnion

To link transcriptional with metabolic activity, we performed short-term (48 h) ^15^NH_4_^+^-labeling experiments in parallel to our metatranscriptomic survey in November 2019. As described above, transcriptional activity of the AOA-LC4 population was high in these samples and exceeded other ammonia oxidizers’ transcriptional levels (Fig. 2). As a measure of metabolic activity, incorporation of ^15^N-ammonium was analyzed at the single-cell level using nanoscale secondary ion mass spectrometry (nanoSIMS) coupled to CARD-FISH identification of AOA cells. All measured AOA cells (*n*_cells_ = 37) were metabolically active and incorporated ammonium at an average rate of 5.04 ± 3.01 amol NH_4_^+^ cell^−1^ d^−1^. The AOA population was significantly more enriched in ^15^N than the remaining picoplankton community (*n*_cells_ = 105, non-parametric Mann–Whitney U-test, *p* < 0.005) (Fig. 4). Using the nanoSIMS results, we calculated single-cell N-based growth rates of 0.012 ± 0.006 d^−1^ for AOA at the *in situ* temperature of 4°C. These rates are one order of magnitude lower than those previously observed in pure cultures of *Nitrosopumilus maritimus*^3^ or for AOA in marine subtropical waters^38^. This is explained by the higher incubation temperature of 28°C in the latter two studies. It should however be noted that two other studies in cold marine waters (ice-water to 4°C)^39,40^ reported AOA growth rates comparable to growth rates of *N. maritimus* at 28°C^3^. The nanoSIMS data was further used to quantify AOA cell dimensions. The average AOA cell in Lake Constance is around 0.54 ± 0.11 μm in length and 0.39 ± 0.10 μm in width and has a volume of 0.048 ± 0.035 μm^3^. This translates into an average biomass of 46 ± 15 fg-C cell^−1^ according to the volume-to-carbon relationship established recently by Khachikyan *et al*.^41^. When taking the total abundance of AOA into account (Fig. 1d), this sums up to 0.5 – 5.5 mg-C m^‒3^ (average 2.0 mg-C m^‒3^) stored in AOA cells in the hypolimnion. In comparison to phytoplankton biomass, which we inferred from depth-integrated Chlorophyll *a* measurements, and considering that the hypolimnion extends over 120 m at our sampling station, this AOA-stored carbon makes up 3.3 – 30.6% (average 12.3%) of carbon stored in phytoplankton over the year’s cycle (Fig. 1e).

**Figure 4:**
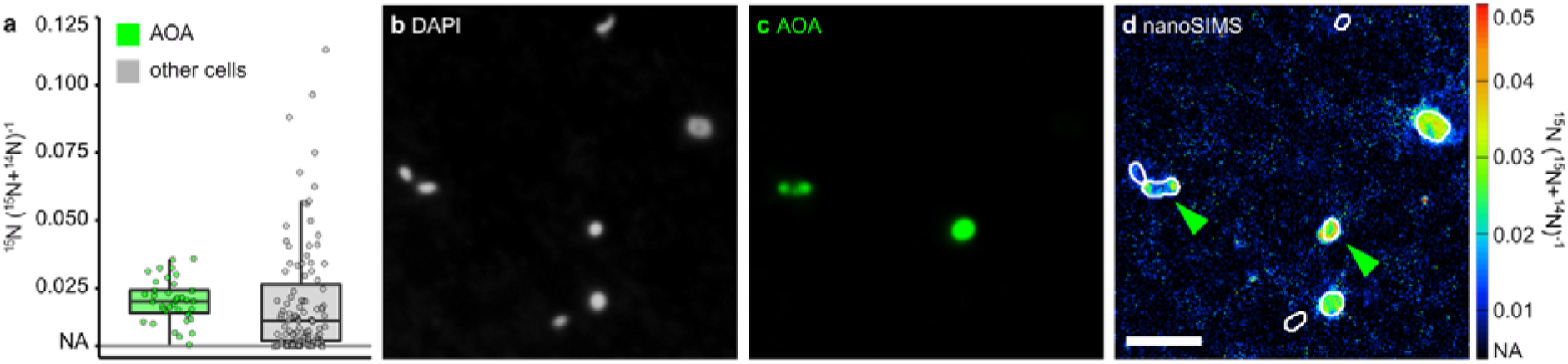
Single-cell ammonium uptake by AOA and non-target picoplankton cells in hypolimnetic water from 85 m depth determined by nanoSIMS combined with CARD-FISH analysis. (a) ^15^N-enrichment of AOA (n=37) and non-target cells (n=105) after addition of ^15^N-ammonium. Representative images of (b) DAPI-stained cells, (c) corresponding CARD-FISH signals with a *Nitrososphaeria*-specific probe, and (d) the corresponding ^15^N-enrichment determined by nanoSIMS; green arrows indicate AOA; scale bar = 2 μm.

### AOA oxidize 1.9 gigagram of ammonium per year in Lake Constance

All independent lines of evidence obtained in this study arrive at the conclusion that the AOA-LC4 population is driving ammonia oxidation in Lake Constance, with a negligible contribution by AOB and comammox bacteria. This natural setting posed the unique opportunity to quantify the ecosystem function exerted by planktonic AOA and to link this information to a single ecotype. We followed nitrification rates in the hypolimnion (85 m depth) by ^15^NH_4_^+^ incubations over four independent time points from June to November 2019. Nitrification rates were very similar within this time period with an average of 6.1 ± 0.7 nmol l^‒1^ d^‒1^ (Table 1). Similar rates were observed in oligotrophic and deep Lake Superior, where total ammonium and nitrate contents were in the range found in Lake Constance^19,42^. Although the obtained rates are potential rates, the absence of an obvious activity lag phase and the linearity of the activities (R^2^ = 0.91 – 1.00) throughout the incubation time (48 h) point towards an active AOA-LC4 population *in situ* and a realistic rate determination (Supplementary Fig. 13). Our previously published estimate of the nitrification rate in Lake Constance was considerably higher but was based on ^15^N-dilution from the nitrate pool^16^ instead of measuring direct conversion of ^15^N-ammonium to ^15^N-nitrite/nitrate (this study). In addition, the previous single time point estimate was based on a high variation of residual ^15^N in replicates. Therefore, we consider the replicated rate measurements presented in this study to be more realistic and conservative.

**Table 1.** Bulk nitrification rates (n=3) and derived cell-specific AOA-driven nitrification rates in the hypolimnion of Lake Constance. Cell-specific nitrification rates were inferred from nitrification rates divided by the respective absolute abundance of AOA measured by archaeal *amoA* qPCR.

| Sampling date<br>(yyyy-mm-dd) | Nitrification rate in the<br>hypolimnion (nmol l<sup>-1</sup> d<sup>-1</sup>) | *amoA* copy numbers<br>(10<sup>4</sup> copies ml<sup>-1</sup>) | Cell-specific nitrification rate<br>(fmol cell<sup>-1</sup> d<sup>-1</sup>) |
| --- | --- | --- | --- |
| 2019-06-18 | 5.89 ± 0.15 | 2.50 ± 0.63 | 0.24 ± 0.06 |
| 2019-07-29 | 4.61 ± 0.41 | n.d. | – |
| 2019-08-28 | 6.06 ± 0.26 | n.d. | – |
| 2019-11-05 | 7.72 ± 0.44 | 3.81 ± 1.55 | 0.20 ± 0.08 |

n.d. = not determined

Measured bulk rates were normalized by archaeal *amoA* copy numbers for the June and November samples to determine cell-specific nitrification rates of the AOA population. At both time points, cell-specific rates were highly similar and averaged to 0.22 ± 0.11 fmol cell^−1^ d^−1^ (Table 1). Like the calculated AOA growth rates, these cell-specific nitrification rates at 4°C are one order of magnitude lower than those observed for cultivated marine *Nitrosopumilus* species at 22 to 28°C^3,43^, or for AOA in marine subtropical waters^44^, as would be expected from the Q_10_ temperature coefficient rule^45^. We used the obtained cell-specific rates to extrapolate AOA-driven nitrification over our complete data set of AOA abundances in Lake Constance (Fig. 1d). As a result, we estimate that about 48 mg NH_4_^+^ m^−3^ y^−1^ is oxidized by the AOA population. This translates to a total of 1.9 gigagram NH_4_^+^ oxidized per year if integrated over the whole lake’s hypolimnion (38.1 km^3^). In comparison, 15.7 gigagram of biomass-N are annually produced by primary production in the photic zone of Lake Constance, when considering the annual primary productivity rate of 220 g C m^−2^ y^−**1** 46^, 473 km^2^ surface area of Lake Constance^24^ and the Redfield ratio (C:N = 106:16). In conclusion, AOA-converted N corresponds to 12% of photosynthetic biomass-N produced in the photic zone of Lake Constance.

### Epilog proposal of “Candidatus Nitrosopumilus limneticus”

Based on its phylogenetic placement, gANI values and habitat preference, we propose a new species name for AOA-LC4: “*Candidatus* Nitrosopumilus limneticus sp. nov.” (lim.ne’ti.cus. N.L. masc. adj. *limneticus* (from Gr. *limne*, lake, swamp) belonging to a lake). “*Candidatus* Nitrosopumilus limneticus” encodes and transcribes genes essential for dissimilatory ammonia oxidation to nitrite and autotrophic CO_2_ fixation via the 3-hydroxypropionyl/4-hydroxybutyryl pathway. Its preferred habitat is the oxygenated hypolimnion of oligotrophic freshwater lakes.

## Conclusion

Over the last two decades, reports accumulated that deep oligotrophic lakes sustain large populations of *Nitrososphaeria* in their hypolimnion^15–20,47^. Our study links these large archaeal populations to nitrification activity and quantifies this important ecosystem service at the single-cell, population, and ecosystem levels. We provide compelling evidence that AOA in a deep oligotrophic lake play an equally important role in the nitrogen cycle—both in terms of their relative abundance and nitrogen fluxes—as their counterparts in marine ecosystems do^21,48^. Lakes are considered sentinels of climate change responding to rising annual temperatures by changes in their physical, chemical and biological properties^49^. Lake Constance is not an exception; its surface waters have become increasingly much warmer, with an annual average increase of 0.9°C observed between 1962 to 2014^50^. Further warming is predicted to continue at 0.03°C y^−1^ resulting in increasing thermal stratification and concomitant increased deoxygenation of deep hypolimnetic waters^50^. Here, we show that nitrification is a key process in the lake’s nitrogen cycle, which we inferred to be driven by a single freshwater AOA ecotype, designated as *Candidatus* Nitrosopumilus limneticus. The results support the general view that the alpha diversity of AOA is extremely low in freshwater lakes^51–54^. In turn, this raises the question how resilient this ecosystem service is to changes in the physical and chemical properties of freshwater lakes in the face of climate change. Answers to this question are interlinked with our quest for clean drinking water supply, since nitrification prevents the accumulation of harmful ammonium, and the quality of freshwater bodies for fisheries and other lacustrine fauna.

## Material and Methods

### Study area and sampling procedure

Lake Constance is a peri-alpine lake with a maximum depth of 251 m, which is monomictic and turning over only in winter. Sampling was conducted at the long-term ecological research station of the University of Konstanz (47.75788° N, 9.12617° E), which is located in the northwestern branch of Upper Lake Constance with a maximum depth of around 140 m (Supplementary Fig. 14). Upper Lake Constance has a permanently oxygenated hypolimnion throughout the year. In this study, we refer to Lake Constance only including deep and oligotrophic Upper Lake Constance and excluding the smaller, shallow and mesotrophic Lower Lake Constance (Supplementary Fig. 14).

Vertical profiles of temperature and oxygen were measured down to the lake sediment with a multi-sampling probe (RBR Ltd.; Ottawa, Canada, Sea & Sun Technology GmbH, Trappenkamp, Germany or bbe Moldaenke GmbH, Schwentinental, Germany). Nitrate and total ammonium (NH_4_^+^ + NH_3_) concentrations were determined using the auto-analyzer and Seal analytics methods G-172-96 Rev. 12 and G-171-96 Rev. 14, respectively (SEAL Analytical GmbH, Norderstedt, Germany). For these measurements, 20 ml water each from 13 depths between 1 − 135 m were taken, filtered through a Chromafil^®^ GF/PET-20/25 filter (pore size 1.0 and 0.2 μm, VWR, Vienna, Austria) and stored at –20°C until analysis. Samples obtained between July and November 2019 were measured by an alternative method: Nitrate was measured by ion chromatography (S150 Chromatography System, SYKAM) and total ammonium was measured fluorometrically by the *ortho*-phthaldialdehyde method^55^. Chlorophyll *a* was sampled from 22 depths over a gradient of 0−60 m and analyzed spectrophotometrically after extraction in hot ethanol as described previously^56^, but without correcting for pheopigments.

For quantitative PCR and metagenome analyses, water was sampled from 85 m depth in three to four replicates (2.5-5.0 l). Water was pre-filtered through a 70 μm and 30 μm nylon mesh (Franz Eckert GmbH, Germany) to remove larger organisms and was filtered on board first through 5 μm and then through 0.2 μm polycarbonate filters (47 mm, Merck, Darmstadt, Germany) using pressurized air. Filters were stored immediately on dry ice on board and at –20°C in the laboratory until further processing. For metatranscriptome analyses, water was sampled in three replicates directly at the desired water depth of 85 m with a WTS-LV *in-situ* pump (McLane research laboratories, Inc., Massachusetts, USA). The pump was programmed to stop running after 1 h, 200 l filtered water or if the flow rate reached a minimum of 500 ml min^−1^ (initial flow rate was set to 3,500 ml min^−1^). The amount of filtered water was recorded. Water was filtered serially through a 30 μm mesh (Franz Eckert GmbH), 5 μm and 0.22 μm filters (142 mm, Merck). Filters with 5 μm and 0.22 μm pore size were stored immediately on dry ice on board and at –80°C in the laboratory until further processing. Sampling of the third replicate of the February and April 2019 samples failed as revealed later on by our inability to extract RNA from these filters despite repeated attempts.

### Nucleic acid extraction, qPCR and CARD-FISH analyses

DNA and RNA were extracted separately from respective 0.22-μm filters after a modified protocol designed previously^57^ as detailed in Supplementary Text. Quantitative PCR assays of total bacterial and archaeal 16S rRNA genes as well as archaeal *amoA* were performed as described recently^16^. The *amoA* qPCR assay was specifically designed to target archaeal *amoA* retrieved from the water column of Lake Constance. Efficiency of qPCR assays targeting the archaeal *amoA* and the total bacterial and archaeal 16S rRNA genes were on average 72.5 ± 4.0% and 91.3 ± 1.6%, respectively. After each run, qPCR specificity was checked with a melting curve. In addition, qPCR products from selected runs were visualized on a 2.5%-agarose gel to verify absence of unspecific PCR products. PCR-inhibitory substances were not evident by qPCR analyses of dilution series of two selected lake water DNA extracts.

For CARD-FISH analysis, 50 ml of lake water was fixed with paraformaldehyde (final concentration 1%, without methanol, Electron Microscopy Sciences) overnight at 4°C. Cells were filtered onto 0.2 μm polycarbonate filters (GTTP, Merck Millipore) and washed with filter-sterilized lake water. Filters were stored at −20°C until analysis. Before CARD-FISH, cells on filter sections were immobilized by embedding in 0.1% low-gelling agarose (MetaPhor™, Lonza, Rockland, ME, USA). CARD-FISH was performed using a HRP-labeled oligonucleotide probe specific for *Nitrososphaeria* (HRP-labeled Thaum726 [GCTTTCATCCCTCACCGTC] and unlabeled competitors [Thaum726_compA: GCTTTCGTCCCTCACCGTC, Thaum726_compB: GCTTTCATCCCTCACTGTC])^58,59^. Negative controls using probe NonEUB [ACTCCTACGGGAGGCAGC]^60^ were performed according to a defined protocol^61^ to exclude unspecific signals. Further CARD-FISH analysis was done as described recently^38^ and in the Supplementary Text.

### Next generation sequencing and bioinformatics processing

Metagenome sequencing libraries were prepared with the NEBNext^®^ Ultra™ DNA Library Prep Kit for Illumina^®^ (New England Biolabs GmbH, Frankfurt am Main, Germany) and sequenced on an Illumina NextSeq500 sequencer using 2 × 150 bp. This resulted in 0.6 – 2.7 × 10^8^ reads per metagenome with an average of 1.3 × 10^8^ reads (8.8 – 41 Gbp, average 20 Gbp). Raw Illumina reads were quality checked with FastQC v.0.11.8^62^ and subsequently quality filtered and trimmed using Sickle v1.33^63^. Thereafter, reads were assembled with Megahit v1.1.2^64^ and binned with maxbin2 v2.2.4^65^. A co-assembly of the Nov 2017, Dec 2017, and Feb 2018 metagenomes resulted in the best AOA bin. To refine this bin, DNA from November 2017 was sequenced in addition by PacBio sequencing on a Sequel instrument (Pacific Biosciences, Menlo Park, CA, USA) using circular consensus sequencing with a target length of 2 kbp. Raw PacBio reads were quality controlled with smrt analysis using an accuracy of 0.999; the number of resulting CCS bases was 0.55 Gbp. The original Illumina AOA bin was used as trusted contigs in spades v3.11.1^66^ and re-assembled with Sequel-reads for gap closure. A subsequent binning in metabat2 v2.12.1^67^ resulted in the final MAG AOA-LC4. MAGs related to AOB, NOB or comammox could not be further refined by long PacBio reads. All MAGs were tested for completeness, strain heterogeneity, and contamination using CheckM v1.0.7^27^ and for their index of replication using iRep v1.10^28^. MAGs and single contigs were screened for the presence of the functional marker genes *amoA* and *nxrB* by both blastp v2.10.1^68^ (e-value threshold 1^−10^) and hmm-search v3.3^69^ (e-value threshold 1^−5^). The latter was based on hmm-models retrieved from the fungene database with manual curation [amoA_AOA.hmm (Feifei Liu), amoA_AOB.hmm (RDP) amoA_comammox.hmm (Yang Ouyang), nxrB.hmm (RDP), fungene.cme.msu.edu]^70^. MAG AOA-LC4 was annotated using the Microscope platform^71^. The automated annotation was manually refined using annotation rules laid out before^72^. Additional MAGs were annotated with PROKKA v1.12^73^ and curated manually for their functional marker genes *amoABC, hao*, and *nxrAB*, where appropriate.

For metatranscriptome sequencing, messenger RNA (mRNA) was enriched from total RNA extracts by depleting ribosomal RNA with the Ribo-off rRNA Depletion Kit for bacteria (Vazyme, Nanjing, China). Thereafter, the sequencing library was prepared with the TruSeq^®^ Stranded mRNA Library Prep (Illumina) and sequenced on a NextSeq500 sequencer using 2 × 150 bp. The sequencing depth ranged between 0.7 – 1.9 × 10^8^ reads per metatranscriptome with an average of 1.2 × 10^8^ reads (10 – 27 Gbp, average 17 Gbp). Raw reads were quality filtered and trimmed using trimmomatic v0.38^74^ and the fastx toolkit v0.0.14 (hannonlab.cshl.edu/fastx_toolkit). Residual ribosomal reads were removed using SortMeRNA v2.1b^75^. Curated metatranscriptome reads were mapped against MAGs and contigs of interest using bowtie2 v2.30^76^ to determine the transcription levels of individual genes. Subsequent network analysis of co-transcribed genes of MAG AOA-LC4 was based on genes with transcription values higher than the median transcription of all AOA-LC4 genes (35.35 FPKM).

Transcription values were correlated pairwise using Spearman correlation; only significant (FDR-corrected *p*-value < 0.05) correlations with a correlation coefficient of *r_S_* > 0.8 were further processed. For the final network construction, only genes, which correlated in their transcriptional response to at least two of the either *amoA*, *amoB* or *amoC* were taken into account. The network was created with the R package igraph v1.2.5^77^ and refined with cytoscape v3.8.1^78^.

### Phylogenetic analyses

Phylogenomic analyses of the nitrifying MAGs were performed on the basis of concatenated amino acid alignments of 122 translated archaeal or 120 bacterial single copy genes^79,80^. Alignments were generated using GTDB-Tk v0.3.2^81^ and maximum likelihood trees were constructed using IQ-tree v1.6.12^82^. Branch support was tested with the Shimodaira–Hasegawa approximate likelihood-ratio test^83^ and ultrafast bootstrap^82^ options in IQ-tree. Genome-wide average nucleic and amino acid identities (ANI and AAI, respectively) were calculated with the online tool (enve-omics.ce.gatech.edu) developed previously^84^ using default settings. Maximum likelihood trees for AOB-*amoA*, comammox-*amoA* and NOB/comammox-*nxrB* genes were constructed in IQ-tree based on manually curated alignments established in ARB v6.0.3^85^.

### Ammonia oxidation rate measurements

To determine ammonia oxidation and ammonium assimilation rates, ^15^N-NH_4_^+^-tracer experiments were carried out as described recently^38^. In June, July, August and November 2019, water was sampled from 85 m depth using a Niskin bottle. Thereafter, 4.5 l were distributed on board to 5-l glass bottles leaving a headspace of 0.5 l. Bottles were sealed with oxalic acid and sodium hydroxide cleaned butyl rubber stoppers and wrapped in aluminum foil to protect them from light. Within 1−7 h after sampling, ^15^N-tracer experiments were started by addition of (^15^NH_4_)_2_SO_4_^2‒^ (10 μM final ^15^N-concentration) and bottles were incubated at *in-situ* temperature (4°C) in the dark. All incubations were done in biological triplicates. Subsamples of 10 ml were taken for nitrification rate measurements over a time period of 48 h (67 h in June), filter-sterilized (0.2 μm) and filters were frozen at −20°C. ^15^N-ammonium labeling percentage was inferred in two steps: First, *in situ* ammonium concentrations were determined using the spectrophotometric salicylate method^86^. Second, *in situ* concentrations were put into relation to added ^15^N-ammonium in samples taken immediately after isotope addition, which was measured by conversion to N_2_ using alkaline hypobromite as described before^87^. ^15^N-ammonium labeling percentage was >99% in all samples. Thereafter, ammonia oxidation rates were determined from combined ^15^N-nitrite and ^15^N-nitrate increase over time. First, ^15^N-nitrite was measured by conversion to N_2_ with sulfamic acid^88^ and ^29^N_2_ was measured by gas chromatography isotope ratio mass spectrometry on a customized TraceGas coupled to a multicollector IsoPrime100 (Isoprime, Manchester, UK). After the ^15^N-NO_2_ measurement, nitrate was reduced to nitrite using spongy cadmium and subsequently converted to N_2_ via sulfamic acid^88,89^. For ammonia oxidation rate calculation, ^15^N-nitrite and ^15^N-nitrate were set to zero at the first sampling time point after tracer addition, and ^15^N-nitrite and ^15^N-nitrate per time point were summed up to give the total ^15^NOx production. Rates were inferred from the slopes of linear regressions; only slopes that were significantly different from zero are reported (P < 0.05, one-sided t-distribution test).

### Single cell ^15^N-uptake measurements from combined CARD-FISH nanoSIMS

After 48 h of the ^15^N-NH_4_^+^-tracer incubation experiment in November 2019, water samples were taken for *Nitrososphaeria*-specific CARD-FISH and subsequent nanoSIMS analyses. A subsample of incubated water (50 ml) was fixed and used for CARD-FISH as described above, but without embedding filter sections in agarose. After CARD-FISH and DAPI staining, regions of interest (ROIs) were marked on a laser microdissection microscope (6000 B, Leica) and images of CARD-FISH signals were acquired on an epifluorescence microscope (Axio Imager, Zeiss). After image acquisition, the filters were sputtered with a 7 nm gold layer to create a conductive surface for nanoSIMS analyses. Single-cell ^15^N-uptake from ^15^N-ammonium was determined using a nanoSIMS 50L (CAMECA), as previously described^90^. Instrument performance was monitored regularly on graphite planchet and on caffeine standards. Due to the small size of AOA (< 1 μm), samples were only briefly (10 − 20 s) pre-sputtered with a Cs+ beam (~300 pA) before measurement. Measurements were performed on a field size of 10 × 10 μm to 15 × 15 μm with a dwelling time of 2 ms per pixel and 256 × 256 pixel resolution over 25 − 40 planes. Analysis of the acquired data was performed using the Look@NanoSIMS software package^91^. Single cell growth and N-assimilation rates were calculated as described recently^44^. We did not account for a possible ^15^N-isotope dilution effect introduced by CARD-FISH^92–94^, as the strength of the effect is dependent on cell growth phase, activity and type, which varies strongly in environmental samples. Therefore, the reported single cell growth and ammonium assimilation rates are a conservative estimate.

### AOA carbon content calculations

The carbon content of individual AOA cells was inferred according to the nonlinear relationship between carbon mass and cell volume (Eq. 1) as recently established by Khachikyan *et al*.^41^:

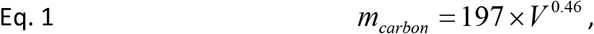

with *m_carbon_* being the mass of carbon in femtograms and *V* being the volume in cubic micrometers of an average AOA cell. *V* was calculated assuming a prolate spheroid cell shape according to Eq. 2 as laid out before^95^:

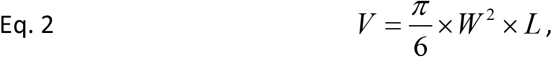

with *W* being the width of the cell and *L* being the length as inferred from the nanoSIMS measurements. Thereafter, the volumetric carbon content of the AOA population was calculated for each time point of our archaeal *amoA* qPCR survey (Fig. 1d) by multiplying the carbon content of a single AOA cell (*m_carbon_*) with the total abundance of AOA (*A_qPCR_*) as inferred by archaeal *amoA* qPCR:

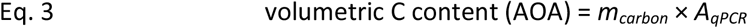

This volumetric C content was integrated over the water column of the hypolimnion and compared to the depth-integrated C content of the phytoplankton, which was inferred from depth-resolved Chlorophyll *a* concentrations using the C : Chl *a* (weight:weight) conversion factor of 31.5 ^96^.

### AOA nitrogen flux calculations

Cell-specific nitrification rates of the AOA-LC4 population (*R_cell-specific_*) were inferred by dividing measured bulk nitrification rates (*R_bulk_*) by the total abundance of AOA as inferred by archaeal *amoA* qPCR (*A_qPCR_*) obtained at the same time points:

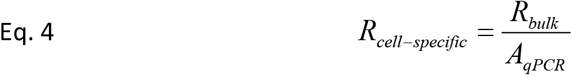

The average of cell-specific nitrification rates was used to infer the annual volumetric nitrification rate of the AOA-LC4 population (*R_annual_*) by integrating over our complete data set of total AOA abundances in the years 2017 to 2019:

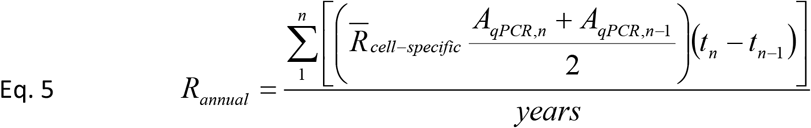

Whenever average values were used to calculate the mean, e.g., to estimate average nitrification rates over the year, the propagation of uncertainty was calculated and the resulting uncertainty provided next to the mean.

## Data Availability

All metagenome and metatranscriptome sequences as well as annotated MAGs are available at NCBI under bioproject number PRJNA691101.

## Acknowledgements

We thank Alfred Sulger as captain of the RV Robert Lauterborn, Joseph Halder, Sylke Wiechmann, Julia Schmidt, Christian Fiek, Isabell Winter, Gabriele Glockgether, Nadine Rujanski, Fenna Alfke, Tine Jordan and Pia Mahler for technical support. We thank Dietmar Straile for provision of chlorophyll *a* data and scientific exchange, Holger Daims for fruitful discussions as well as David Schleheck for providing laboratory space. This research was funded by the German Research Foundation (DFG, GRK 2272/1, project C2 to M.P. and B.S.), the University of Konstanz, the Leibniz Institute DSMZ and the Max Planck Society.

## Author contributions

F.K., K.K., B.S., M.M.M.K. and M.P. designed the study; P.B. performed sequencing and sequencing library preparations and measured total ammonium; K.K. and S.L. ran nanoSIMS analyses; F.K., K.K., D.K.N. and M.P. analyzed samples and data. The manuscript was written by F.K. and M.P. with contributions from all co-authors.

## Additional information

Supplementary information is available for this paper.

