## Supplementary Information for "Quantification of archaea-driven freshwater nitrification: from single cell to ecosystem level"

### Supplementary Text

*Methods: DNA and RNA extraction*

For DNA extraction, 0.22 µm-filters (47 mm diameter) were placed in 2 ml-screw cap tubes and vortexed for 15 min in a solution containing 500 µl TE-buffer (10 mM Tris-HCl pH 8.0, 1 mM EDTA), 12.5 µl 20% sodium lauryl sulfate-solution (SLS; Sigma-Aldrich, Taufkirchen, Germany), 500 µl phenol-chloroform-isoamylalcohol 25:24:1 (Carl Roth GmbH), and 200 µl pre-combusted zirconium beads (0.1 mm in diameter, Carl Roth GmbH, Karlsruhe, Germany). After centrifugation (4°C, 10 min, 18,620 ×*g*), the supernatant was washed once with 500 µl chloroform-isoamylalcohol 24:1 (Carl Roth GmbH). This mixture was centrifuged again and DNA precipitated from the separated supernatant over night at −20°C using a mixture of 0.1 volume 3 M sodium acetate (pH 4.8), 2.5 volume molecular grade ethanol (Carl Roth GmbH) and 1 µl glycogen (20 mg $ml^{-1}$, Thermo Fisher Scientific, Waltham, MA, USA). Precipitated DNA was centrifuged, washed twice with 80% molecular grade ethanol, and dissolved in nuclease-free water (MP biomedicals, Eschwege, Germany). RNA was removed by an RNase ONE™ ribonuclease treatment following kit instructions (Promega, Fitchburg, WI, USA) and DNA samples were stored at –20°C until processing.

For RNA extraction, 0.22 µm-filters (142 mm diameter) filters were cut with a sterilized scissor into thirds and extracted as described above except for the following modifications: filters were extracted in extraction buffer (50 mM sodium acetate and 10 mM EDTA, pH 4.2) with 0.025% SLS (Sigma-Aldrich) and phenol-chloroform-isoamylalcohol 25:24:1 (Roti-Aqua-P/C/I 4.5-5.0, Carl Roth GmbH).

Washing of the aqueous phase with chloroform-isoamylalcohol 24:1 was done in the presence of 0.1 volume 3 M sodium acetate. RNA was finally precipitated with 1 volume ice-cold isopropanol in the presence of 1 µl glycogen (35 mg $ml^{-1}$, RNase-free, VWR), washed as stated above, and eluted in nuclease-free water (MP biomedicals). DNA was digested with the TURBO DNA-free™ kit (Thermo Fisher Scientific) and RNA samples were stored afterwards at –80°C until sequencing.

*Methods: Determination of AOA abundance by CARD-FISH*

Before CARD-FISH, cells on the filter sections were immobilized by embedding in 0.1% low-gelling agarose (Metaphor). CARD-FISH was performed using a specific HRP-labeled oligonucleotide probe for *Nitrososphaeria* (HRP-labeled Thaum726 [GCTTTCATCCCTCACCGTC] and unlabeled competitors [Thaum726_compA: GCTTTCGTCCCTCACCGTC, Thaum726_compB: GCTTTCATCCCTCACTGTC])[1,2] as described previously[3]. Negative controls using NonEUB[4] to exclude unspecific signals were performed according to a defined protocol[5]. Briefly, endogenous peroxidases were inactivated by incubation in 0.01 M HCl for 10 min. Cells were permeabilized by HCl (0.1 M HCl for 1 min) and subsequently washed with MilliQ water. Filter pieces were hybridized with HRP probes and the respective competitor probes at 25% formamide concentration at 46°C for up to 3 h. After a 5 min washing step at 48°C and HRP probe equilibration in 1× PBS for 5 to 15 min, signal amplification was performed with OregonGreen488-labeled tyramides at 48°C for 30 min. Cells were counterstained with 4′,6-diamidino-2-phenylindole (DAPI, 10 µg $ml^{-1}$, 5 min at room temperature). Filter sections were mounted onto glass slides, and embedded in a 4:1 mixture of Citifluor AF1 and Vectashield (Citifluor Ltd, London, UK; Vector Laboratories, Burlingame, CA, USA). *Nitrososphaeria* and DAPI signals were counted on an Axiophot or Axioplan 2 microscope (Zeiss, Germany).

*Results: Phylogenetic analysis of bacterial ammonia oxidizers*

Phylogenomic maximum likelihood tree construction revealed that MAG AOB-LC263 formed a stable cluster with other freshwater MAGs, which represented a sister clade to *bona fide Nitrosospira* species (Supplementary Fig. 5). This was corroborated by phylogenetic analysis of its *amoA* gene (Supplementary Fig. 7). Closest relatives of AOB-LC263 were MAGs retrieved from Lake Baikal, the Great Lakes and Lake Biwa. Based on the currently proposed species and genus delineation thresholds of ca. <95% ANI and <65% AAI, respectively[6], AOB-LC263 would represent a new species and genus within the *Nitrosomonadaceae* (Supplementary Fig. 6). The phylogenetic affiliation of contigs AOB-LC199628 and AOB-LC368213 could only be assessed based on their *amoA* genes. Contig AOB-LC199628 clustered in a stable clade consisting of environmental sequences that was distinct from the AOB-LC263 and *Nitrosospira* clusters. Its closest cultured relative was *Nitrosospira* sp. Np39-19 as based on 78.3% *amoA* nucleotide identity. Contig AOB-LC368213 clustered within sequences affiliated with *Nitrosomonas* species with its closest cultured relative being *Nitrosomonas ureae*

Nm10 with 89.9% *amoA* nucleotide identity (Supplementary Fig. 7). Comparison of the retrieved *amoA* sequences to a previous bacterial *amoA* clone library obtained from Lake Constance waters[7] revealed that the *amoA* gene of contig AOB-LC199628 was 100% identical (nucleic acid identity) to clone BmcYyy23.2 (MH780622.1) from OTU1[7]. Furthermore, the *amoA* of MAG AOB-LC263 was 99.8% identical to clone BmcYyy33 (MH780602.1) from OTU2. Since only two OTUs were detected previously[7], contig AOB-LC368213 had no representatives in the earlier *amoA* clone library.

*Results: Phylogenetic analysis of nitrite-oxidizing bacteria and comammox bacteria*

Phylogenomic maximum likelihood tree construction placed the two MAGs NOB-LC29 and NOB-LC32 into *Nitrospira* lineage II but outside the intra-lineage comammox clades A and B (Supplementary Fig. 8). This was corroborated by phylogenetic analysis of their *nxrB* genes (Supplementary Fig. 10). The two MAGs shared an ANI and AAI of 85% and 85%, respectively, indicating that they represent two separate species within the genus *Nitrospira* (Supplementary Fig. 9). Phylogenomic tree construction of MAG COM-LC224 placed it into comammox clade B within *Nitrospira* lineage II (Supplementary Fig. 8), which was corroborated by phylogenetic placement of its single genes *nxrB* and *amoA* (Supplementary Fig. 10 and 11). Interestingly, COM-LC224 showed <65% AAI to both type and *Candidatus* species within the genus *Nitrospira* (Supplementary Fig. 9), but at the same time exhibited AAI values of >65% with freshwater *Nitrospira* like NOB-LC29 and NOB–LC32. Without further data, its affiliation at the taxonomic rank of a genus is currently inconclusive.

### Supplementary Tables

**Supplementary Table 1.** Overview of metagenome-assembled genomes (MAGs) and contigs related to the nitrifying community in the hypolimnion of Lake Constance.

**Supplementary Table 2.** Annotation and seasonally resolved transcription of MAG AOA-LC4 genes involved in nitrogen metabolism, vitamin synthesis, carbon fixation, cell division, replication, transport systems, respiration and the TCA-cycle.

**Supplementary Table 3.** NCBI accession numbers or Taxon IDs of all species, MAGs and clones, which are part of the phylogenetic trees of *Nitrososphaeria*, *Nitrosomonadaceae* and *Nitrospira* (Supplementary Figures 2, 5, 7, 8, 10, and 11).

### Supplementary Figures

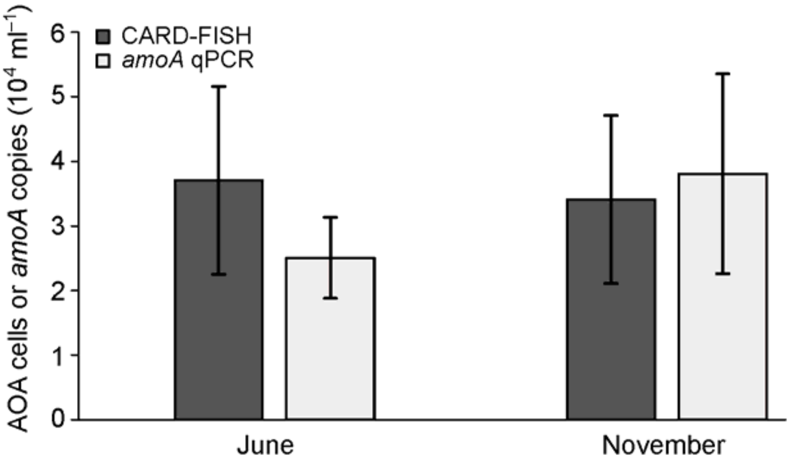


**Supplementary Figure 1.** AOA abundance in hypolimnetic water from 85 m depth as measured by archaeal *amoA*-targeted qPCR or CARD-FISH using a *Nitrososphaeria*-specific probe (probe Thaum726), which currently encompasses all AOA. Samples were taken on June 18th and November 5th 2019. CARD-FISH was performed on water samples used for nitrification rate measurements after 67 h (June) or 48 h (November) of incubation at 4°C in the dark. Hybridized filters were counted either 10 times (June) or 25 times (November) independently.

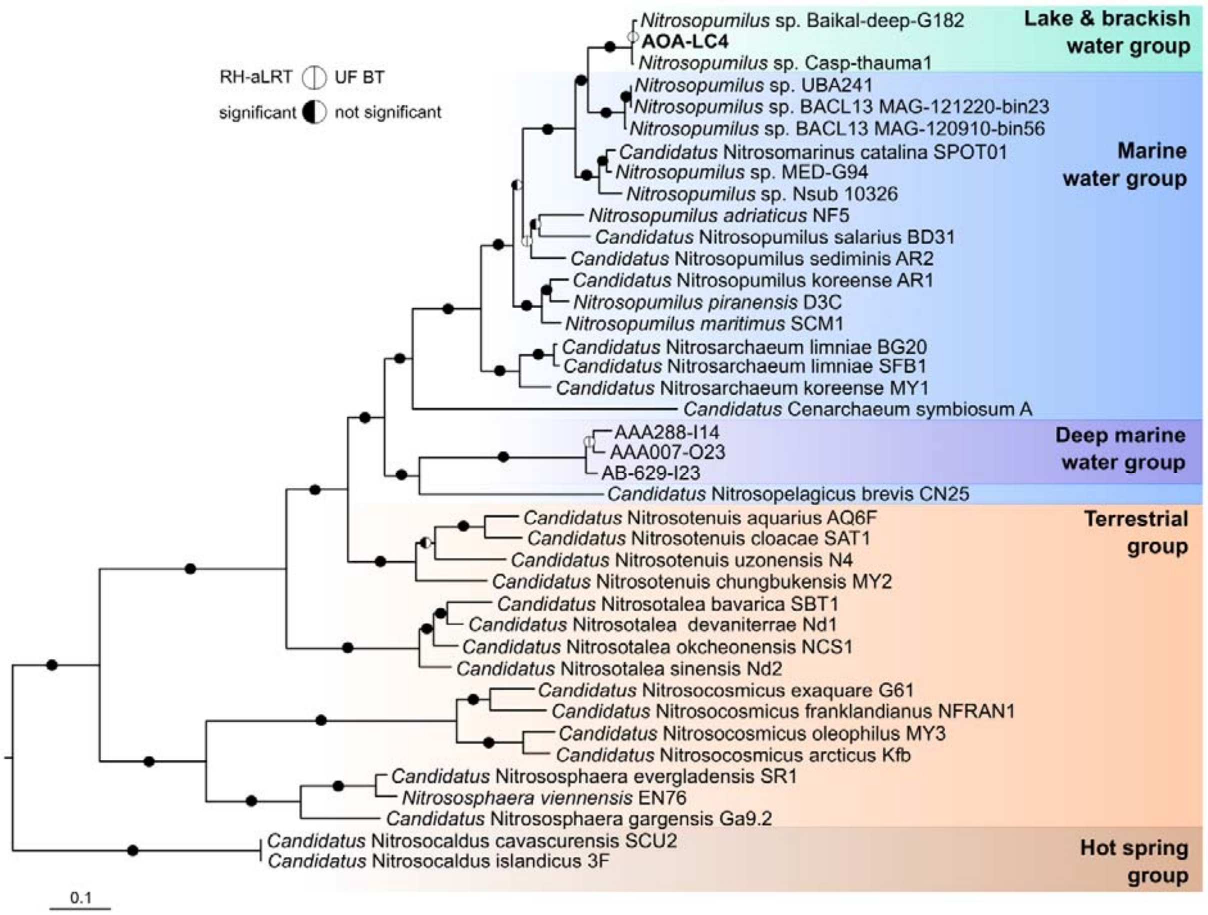


**Supplementary Figure 2.** Phylogeny of MAG AOA-LC4 in relation to MAGs retrieved from other major inland water bodies within the class *Nitrososphaeria*. The phylogenomic maximum likelihood tree was re-constructed using the IQ-tree algorithm[8] on the basis of a concatenated amino acid alignment of 122 translated single-copy genes that were established by the GTDB-based taxonomy for phylogenetic inference of archaea[9]. Branch support was tested with the Shimodaira–Hasegawa approximate likelihood-ratio test (SH-aLRT; 1000 replicates) and ultrafast bootstrap (1000 replicates) using the IQ-tree software package[8]. Branch support was set as significant at ≥80% for SH-aLRT and ≥95% for ultrafast bootstrap values (black semi-circles for significant and white for non-significant). Non-AOA Thaumarchaeota archaeon sp. BS3 (IMG Taxon ID 2721755844), unclassified Thaumarchaeota DRTY-7 bin 36 (2263082001), and Thaumarchaeota archaeon strain DS1 (2263082000) were used as outgroup. The scale bar indicates 10% estimated amino acid sequence divergence. Accession numbers or respective Taxon IDs can be found in Supplementary Table 3.

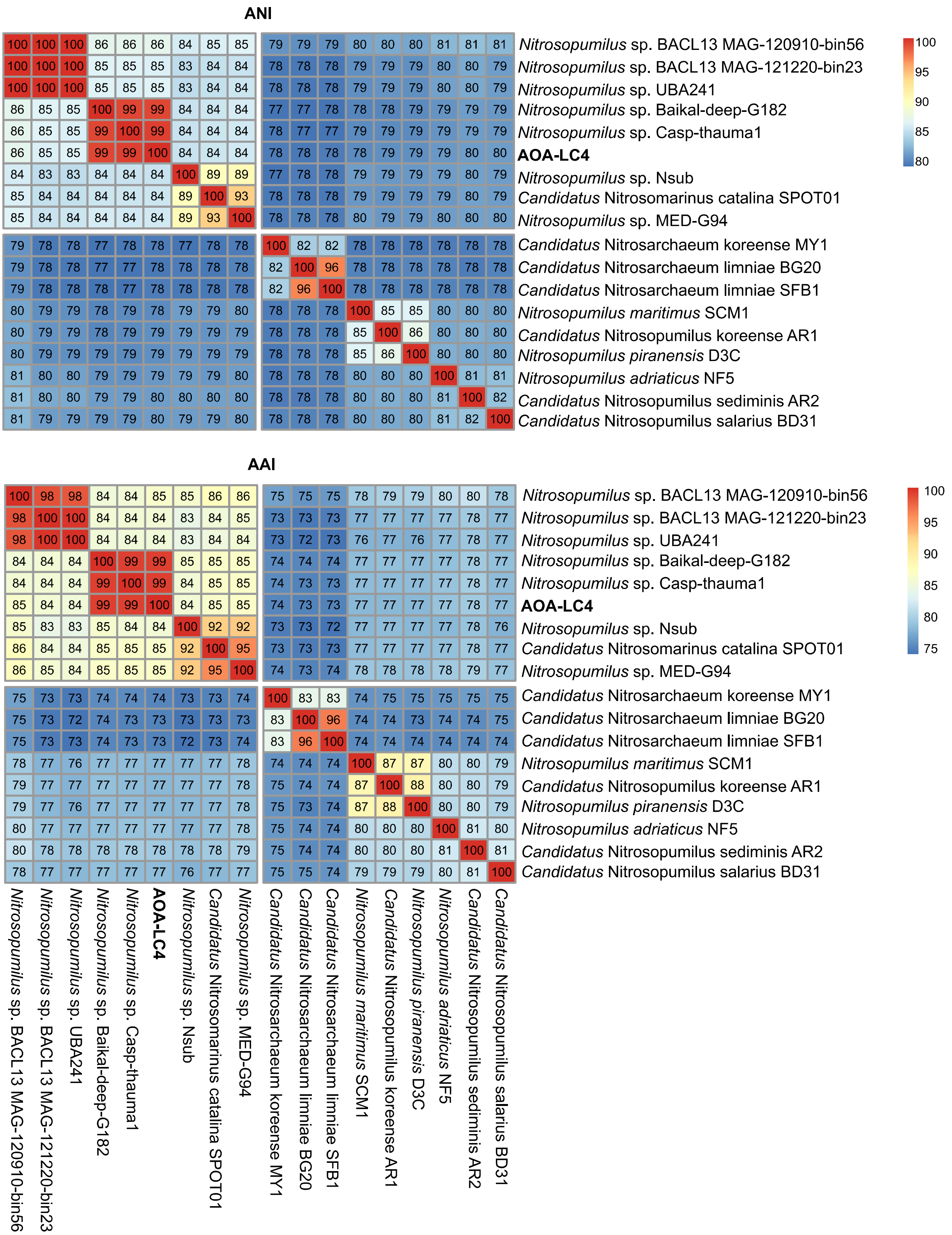


**Supplementary Figure 3.** Pairwise genome-wide average nucleotide identities (ANI) and average amino acid identities (AAI) of MAG AOA-LC4 (shown in bold) in comparison to representatives of the family *Nitrosopumilaceae*. MAG AOA-LC4 represents a novel species together with Casp-thauma1 and Baikal-Deep-G182 in the genus *Nitrosopumilus* based on the species-level threshold of 95% for ANI[6] and genus-threshold of 65% for AAI[6].

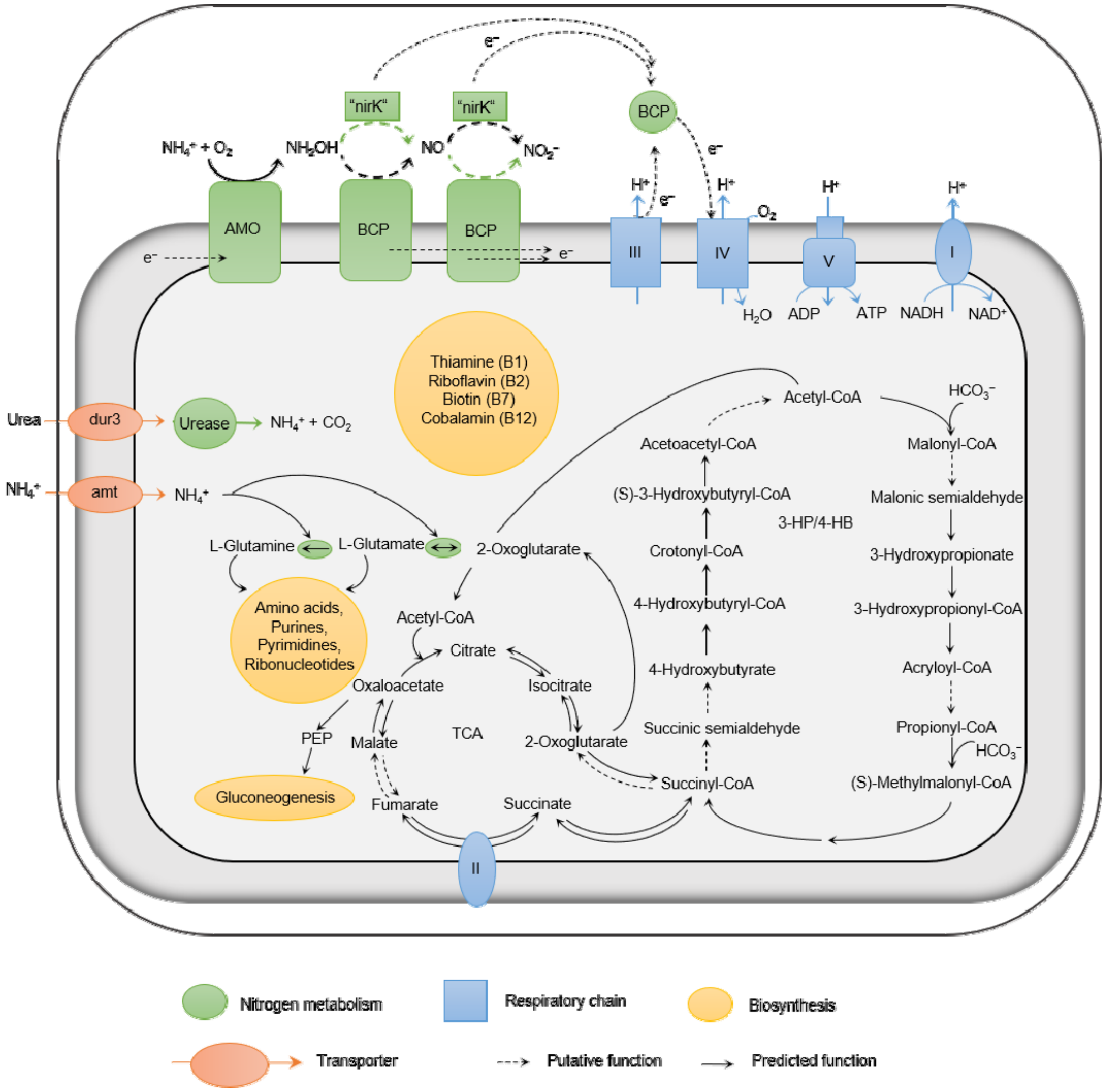


**Supplementary Figure 4.** Predicted metabolism of MAG AOA-LC4. The two proposed alternative pathways for ammonia oxidation in AOA were taken from models published previously[10,11] and are indicated by either green or black dashed arrows. Abbreviations: AMO: ammonia monooxygenase, "NirK": multicopper oxidase with putative nitrite reductase activity, BCP: putative blue (type 1) copper domain protein, 3-HP/4-HB: 3-hydroxypropionyl/4-hydroxybutyryl pathway, TCA: tricarboxylic acid cycle. All genes encoding presented pathways are listed in Supplementary Table 2 with further annotation information and seasonally resolved transcription values at 85 m depth.

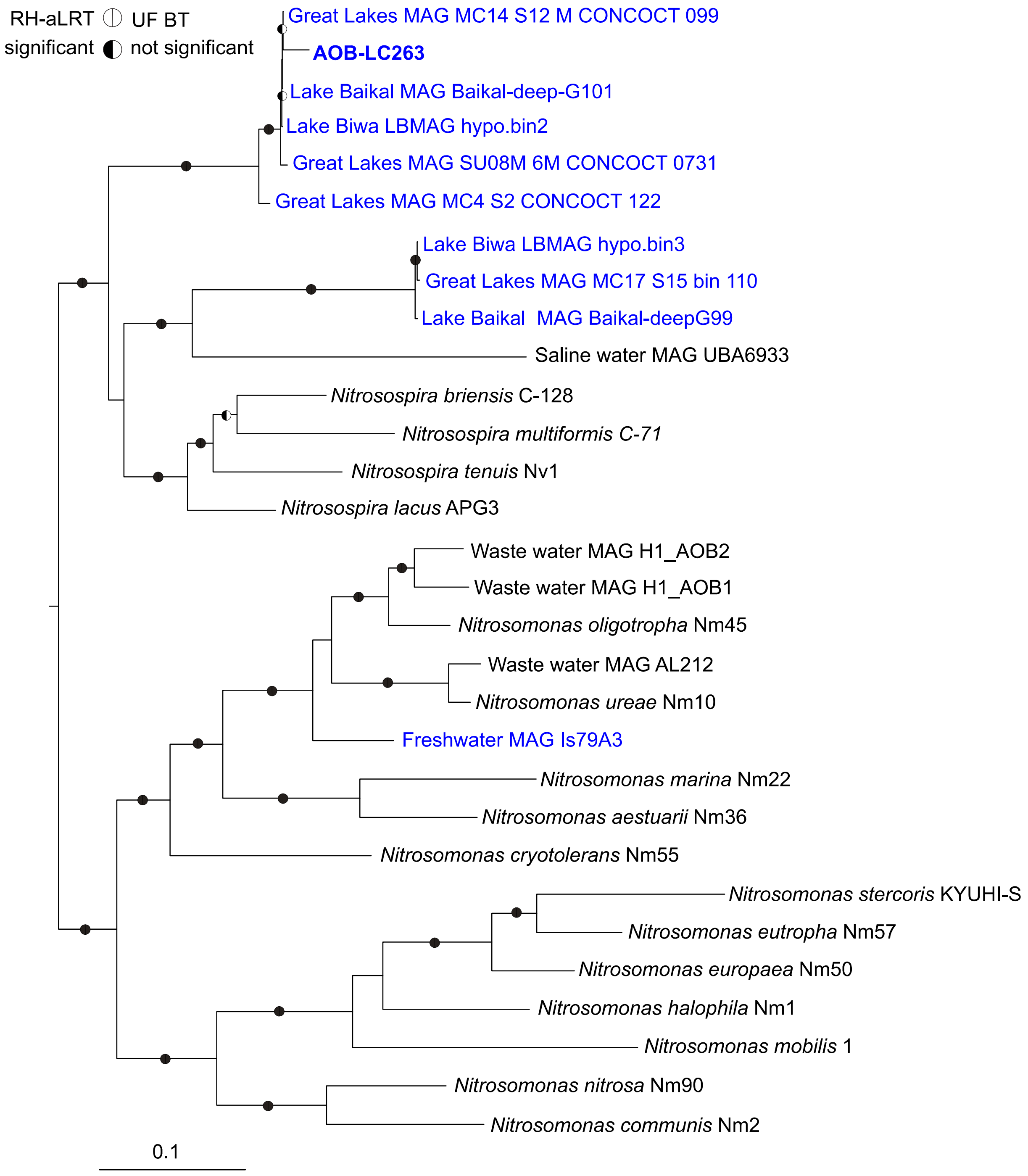


**Supplementary Figure 5.** Phylogeny of MAG AOB-LC263 in relation to closely related freshwater MAGs and pure cultures of the ammonia-oxidizing bacteria within the genera *Nitrosomonas* and *Nitrosospira*. The phylogenomic maximum likelihood tree was constructed using the IQ-tree algorithm[8] on the basis of a concatenated amino acid alignment of 120 translated single copy genes that were established by the GTDB-based taxonomy for phylogenetic inference of bacteria[12,13]. Branch support was tested with the Shimodaira–Hasegawa approximate likelihood-ratio test (SH-aLRT; 1000 replicates) and ultrafast bootstraps (1000 replicates) within IQ-tree. Branch support was set as significant at ≥80% for SH-aLRT and ≥95% for ultrafast bootstrap values (black semi-circles for significant and white for non-significant). MAGs or species with freshwater-origin are colored blue.

*Methylotenera mobilis* (NCBI accession number GCA_000023705.1), *Methylovorus glucosetrophus* (NC_012969.1) and *Methylobacillus flagellates* (GCA_000013705.1) were used as outgroup. The scale bar indicates 10% estimated amino acid sequence divergence. All accession numbers can be found in Supplementary Table 3.

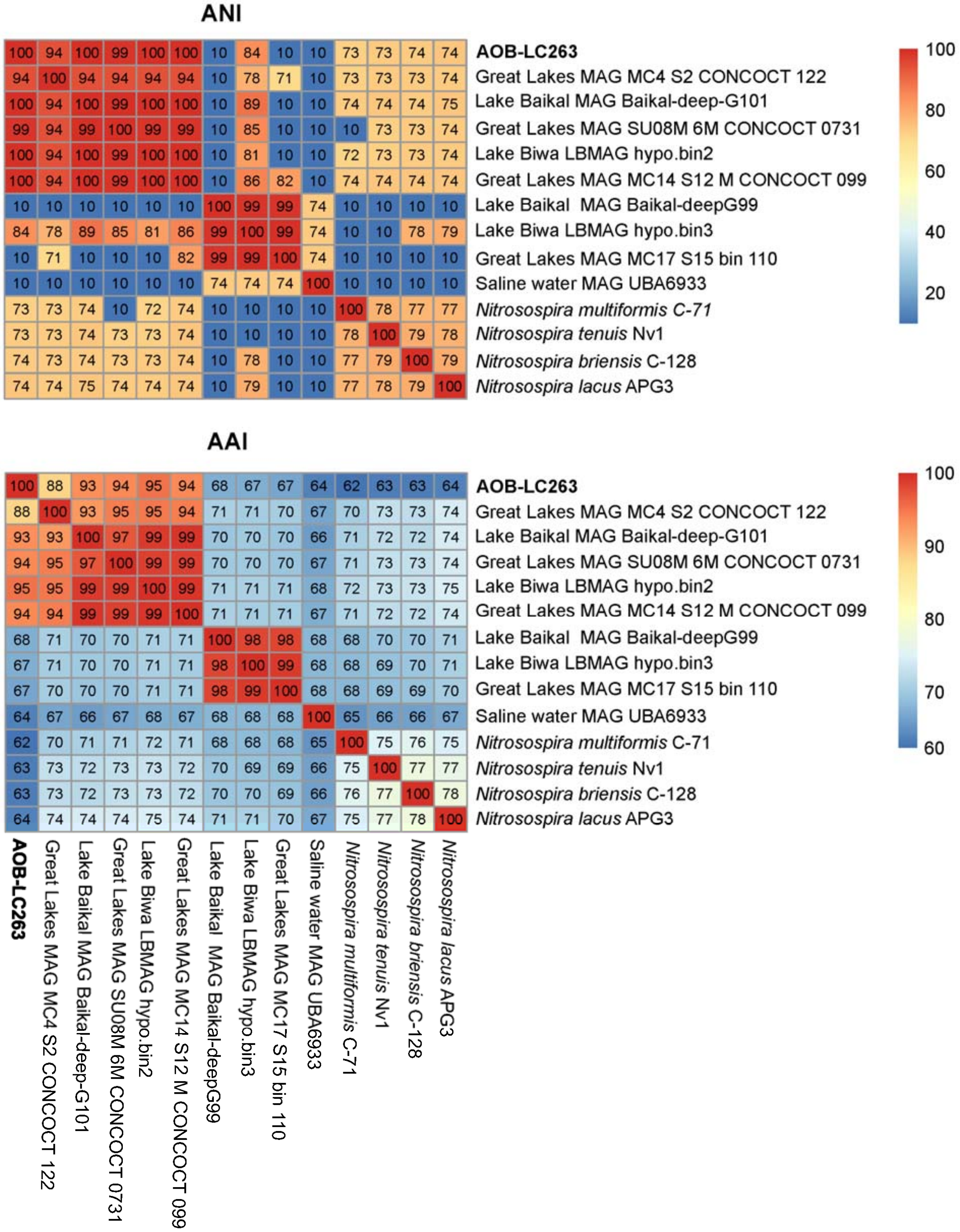


**Supplementary Figure 6.** Pairwise genome-wide average nucleotide identities (ANI) and average amino acid (AAI) identities for MAG AOB-LC263 (shown in bold) in comparison to closely related MAGs and representatives of the genus *Nitrosospira*. AOB-LC263 represents a novel genus compared to described *Nitrosospira* species based on the threshold of 65% for AAI[6].

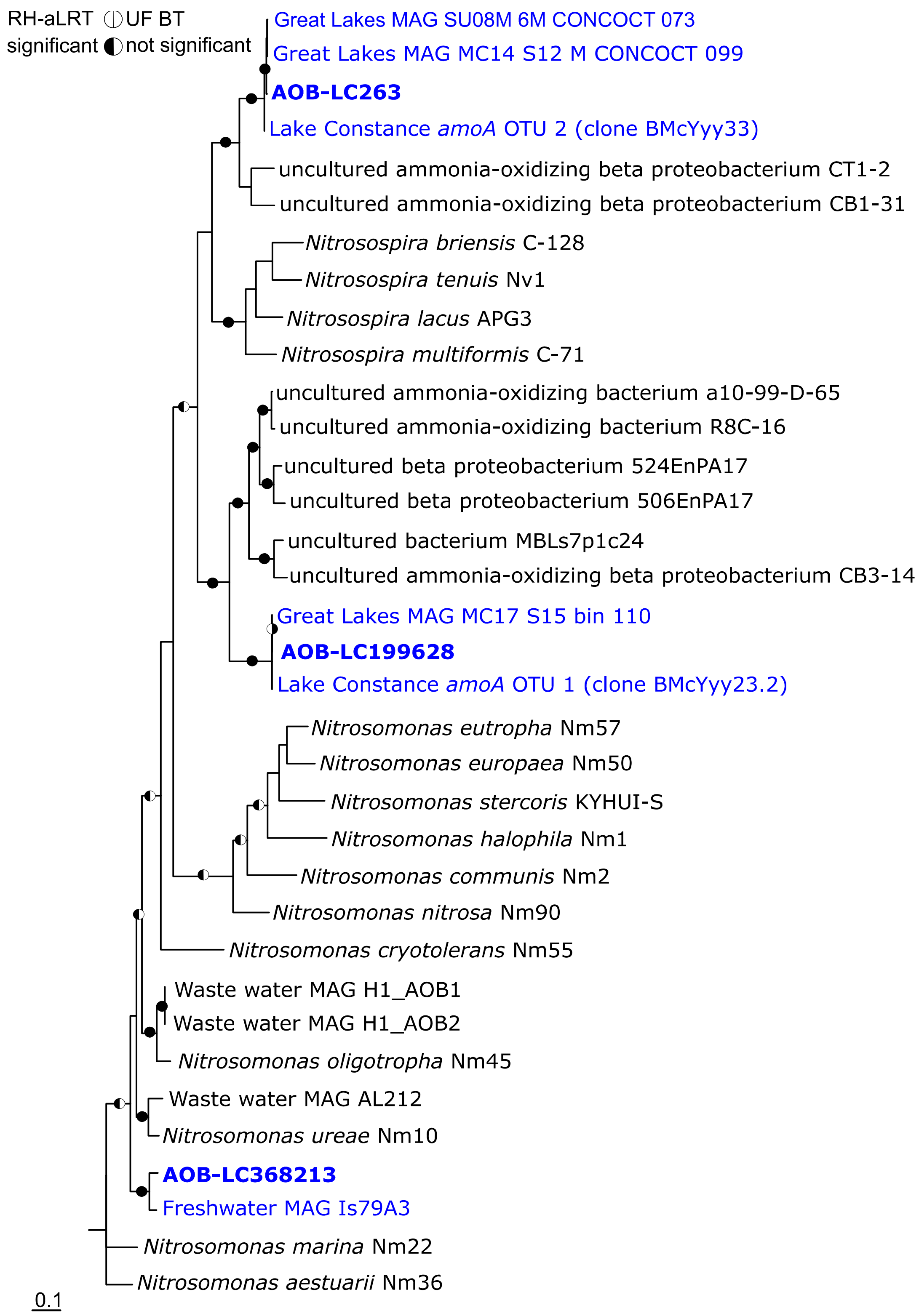


**Supplementary Figure 7.** Phylogeny of MAG AOB-LC263 and contigs AOB-LC199628 and AOB-LC368213 in relation to ammonia-oxidizing bacteria and environmental sequences affiliated with the family *Nitrosomonadaceae* as based on the functional marker gene *amoA*. The maximum likelihood tree was inferred by the IQ-tree algorithm[8] using 459 unambiguous alignment positions of the bacterial *amoA* gene. Branch support was tested with the Shimodaira–Hasegawa approximate likelihood-ratio test (SH-aLRT; 1000 replicates) and ultrafast bootstraps (1000 replicates) within IQ-tree. Branch support was set as significant at ≥80% for SH-aLRT and ≥95% for ultrafast bootstrap values (black semi-circles for significant and white for non-significant). MAGs, clones or species with freshwater-origin are colored blue. *Nitrosococcus watsonii* (NC_014315) and *Nitrosococcus oceani* (NC_007484) *amoA* genes were used as outgroup. The scale bar indicates 10% estimated nucleic acid sequence divergence. Accession numbers can be found in Supplementary Table 3.

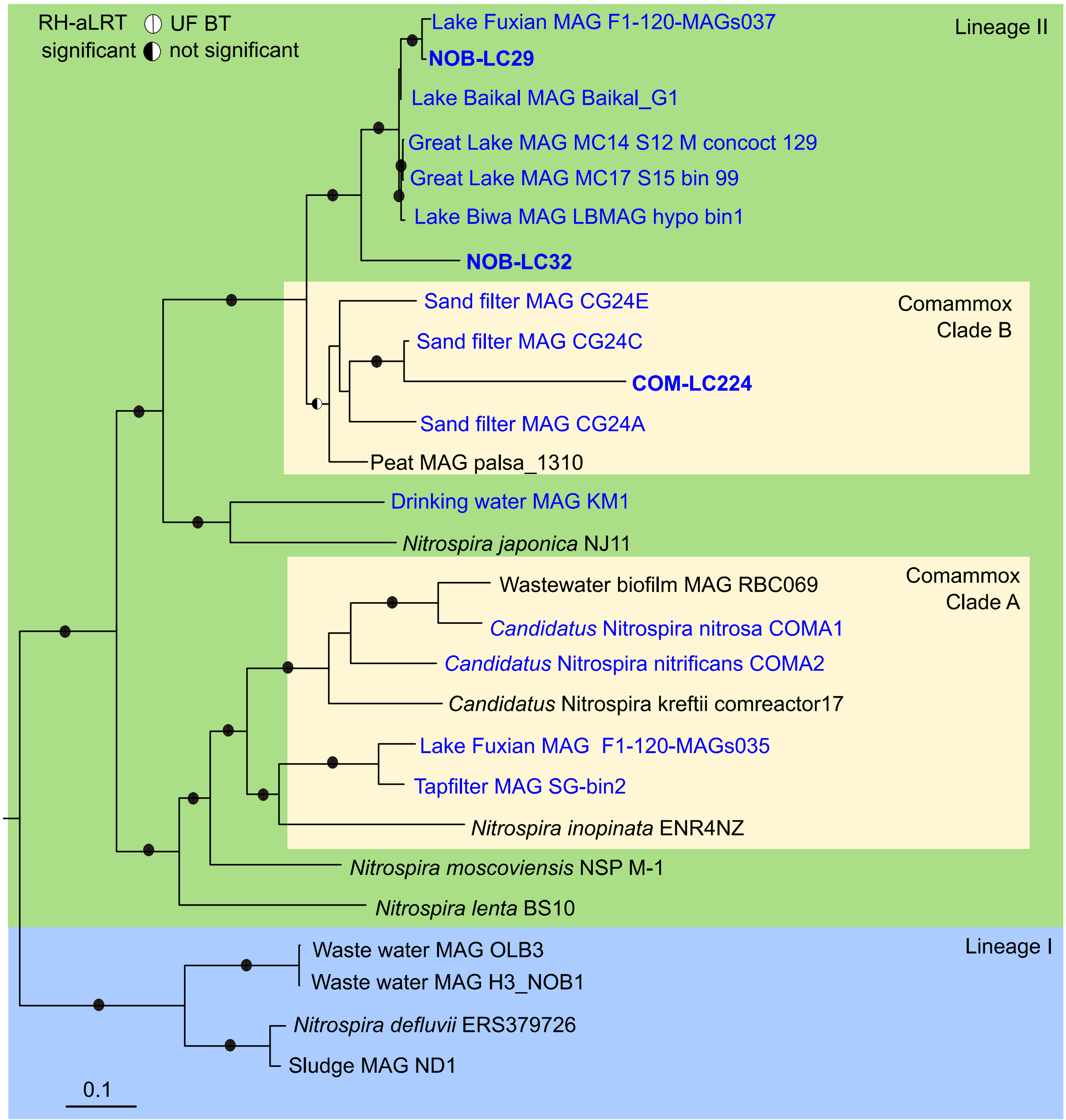


**Supplementary Figure 8.** Phylogeny of MAGs NOB-LC29, NOB-LC32, and COM-LC224 in relation to representatives of *Nitrospira* lineage I and II. MAGs affiliated with either comammox clade A or B were taken from the literature[14,15]. Clade classification of comammox bacteria is based on their *amoA* gene as proposed by Daims *et al.*[16] and Pjevac *et al.*[17]. The phylogenomic maximum likelihood tree was constructed using the IQ-Tree algorithm[8] on the basis of a concatenated amino acid alignment of 120 translated single copy genes that were established by the GTDB-based taxonomy for phylogenetic inference of bacteria[12,13]. Branch support was tested with the Shimodaira–Hasegawa approximate likelihood-ratio test (SH-aLRT; 1000 replicates) and ultrafast bootstraps (1000 replicates) within IQ-tree. Branch support was set as significant at ≥80% for SH-aLRT and ≥95% for ultrafast bootstrap values (black semi-circles for significant and white for non-significant). MAGs or species with freshwater-origin are colored blue. *Leptospirillum ferriphilum* (GCA_900198525.1) and *Leptospirillum ferrooxidans* (GCA_000284315.1) were used as outgroup. The scale bar indicates 10% estimated amino acid sequence divergence. Accession numbers can be found in Supplementary Table 3.

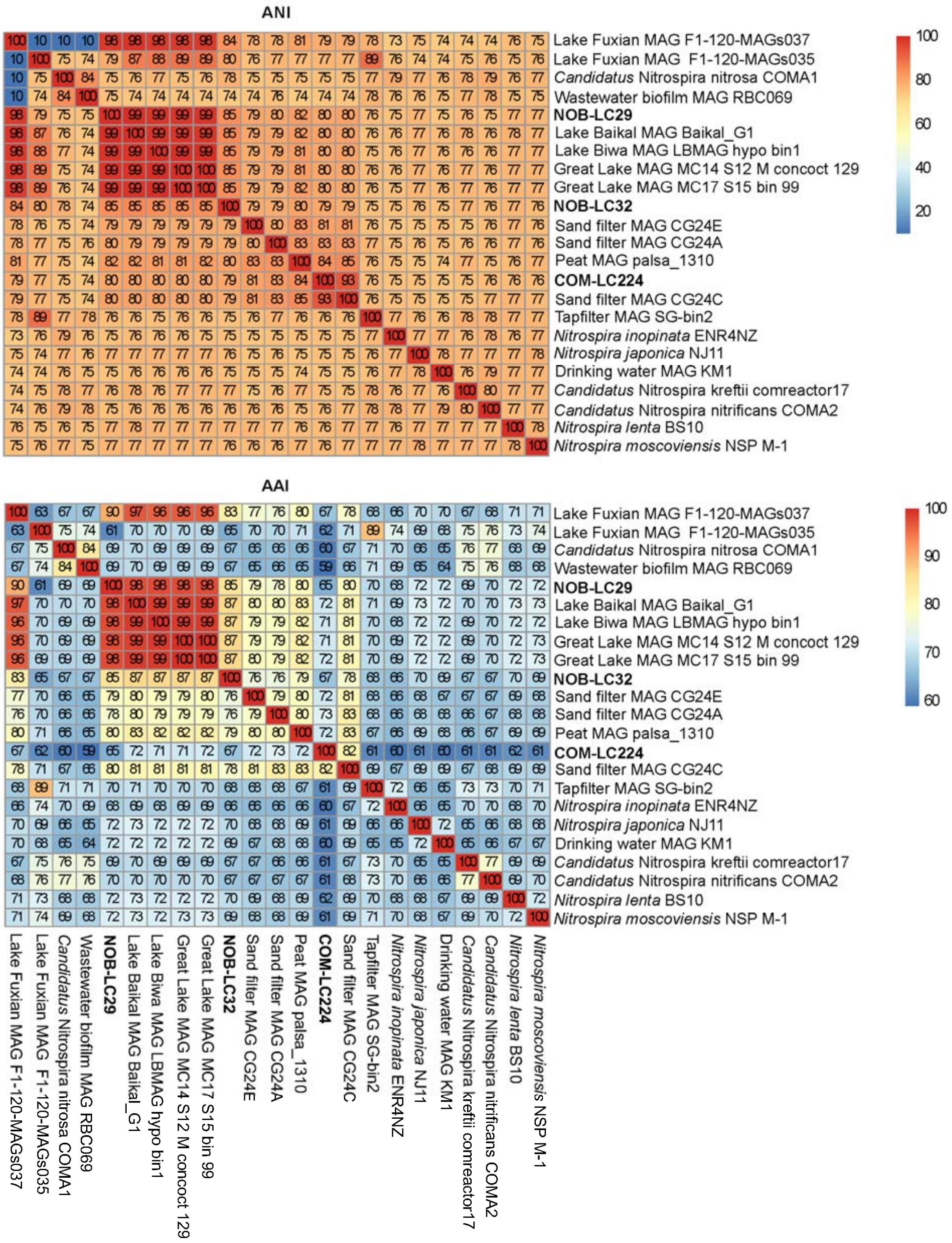


**Supplementary Figure 9.** Pairwise genome-wide average nucleotide identities (ANI) and average amino acid identities (AAI) for lineage II *Nitrospira* including MAGs NOB-LC32, NOB-LC29, and COM-LC224 (shown in bold).

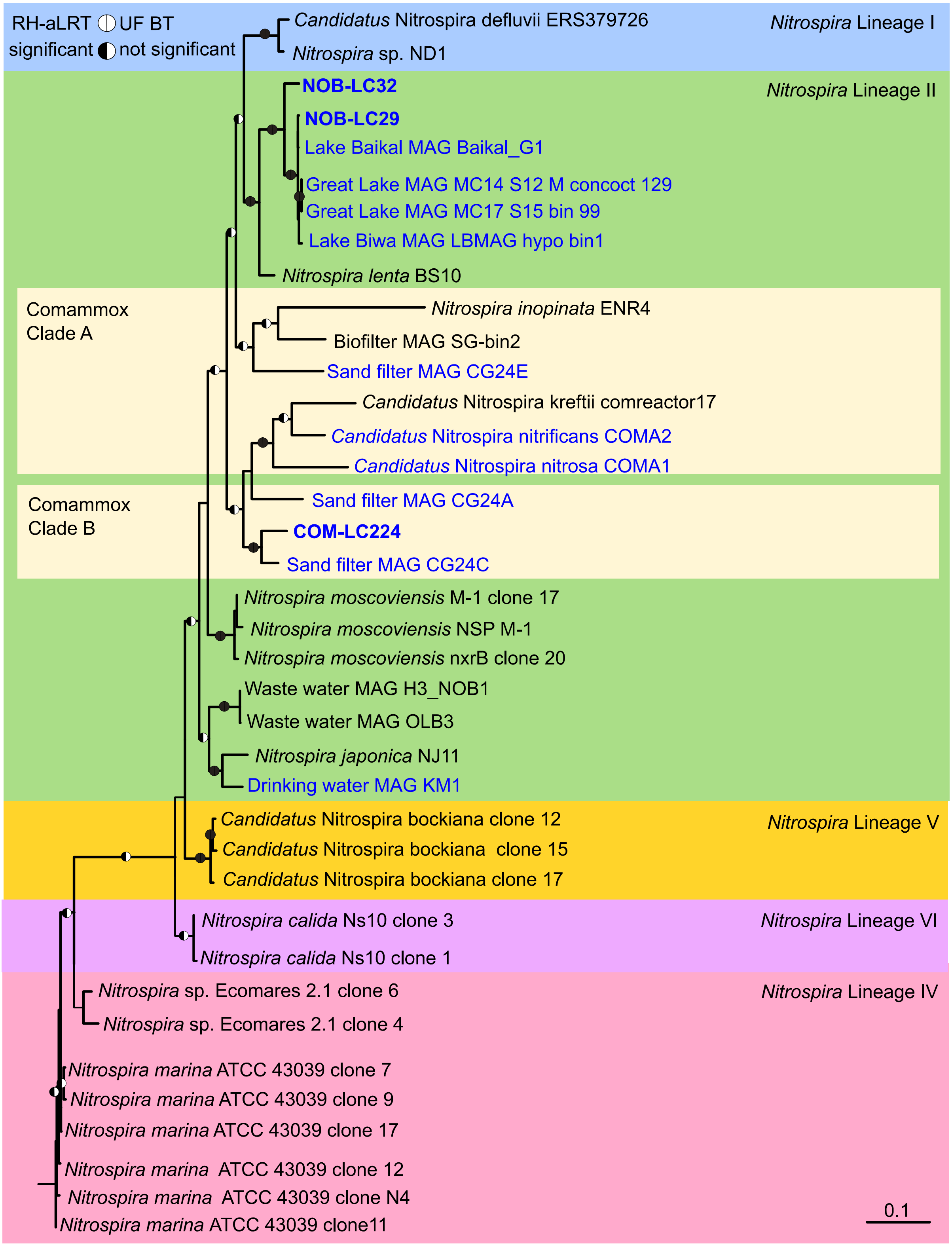


**Supplementary Figure 10.** Phylogeny of MAGs NOB-LC29, NOB-LC32, and COM-LC224 as based on the functional marker gene *nxrB*. Classification of the comammox MAGs into clade A or B was transferred from the phylogenomic tree (Supplementary Fig. 8), as comammox species cannot be distinguished based on their *nxrB* gene from canonical *Nitrospira* species[16,18]. The maximum likelihood tree was inferred by the IQ-tree algorithm[8] using 1,205 unambiguous alignment positions of the *nxrB* gene. Branch support was tested with the Shimodaira–Hasegawa approximate likelihood-ratio test (SH-aLRT; 1000 replicates) and ultrafast bootstrap (1000 replicates) within the IQ-tree software package. Branch support was set as significant at ≥80% for SH-aLRT and ≥95% for ultrafast bootstrap values (black semi-circles for significant and white for non-significant). MAGs, clones or species with freshwater-origin are colored blue. *Hydrogenobaculum* sp. (NC_011126), *Natronomonas pharaonis* (NC_007426.1) and *Candidatus* Kuenenia stuttgartiensis (CT573072) *narH*-like genes were used as outgroup. The scale bar indicates 10% estimated nucleic acid sequence divergence. Accession numbers can be found in Supplementary Table 3.

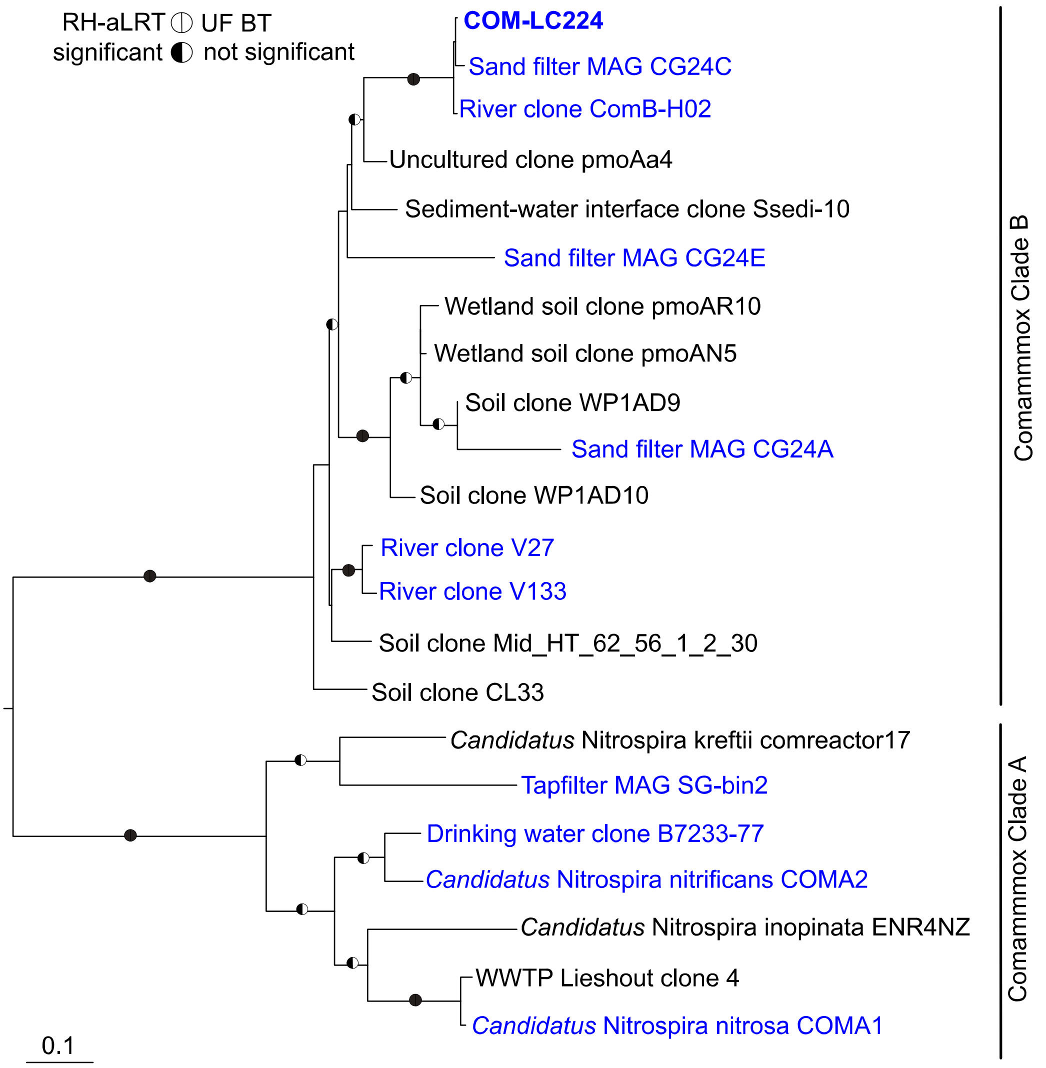


**Supplementary Figure 11.** Phylogeny of COM-LC224 as based on the functional marker gene *amoA*. Classification into clade A or B was done as proposed by by Daims *et al.*[16] and Pjevac *et al*.[17]. The maximum likelihood tree was inferred by the IQ-tree algorithm[8] using 414 unambiguous alignment positions of the *amoA* gene. Branch support was tested with the Shimodaira–Hasegawa approximate likelihood-ratio test (SH-aLRT; 1000 replicates) and ultrafast bootstrap (1000 replicates) within the IQ-tree software package. Branch support was set as significant at ≥80% for SH-aLRT and ≥95% for ultrafast bootstrap values (black semi-circles for significant and white for non-significant). MAGs or species with freshwater-origin are colored blue. *Nitrosospira multiformis* (U91603), *Nitrosospira* sp. (WP_041514847.1*),* *Nitrosomonas europaea* (L08050) and *Nitrosomonas communis* (WP_046851395) *amoA* genes were used as outgroup. The scale bar indicates 10% estimated nucleic acid sequence divergence. Accession numbers can be found in Supplementary Table 3.

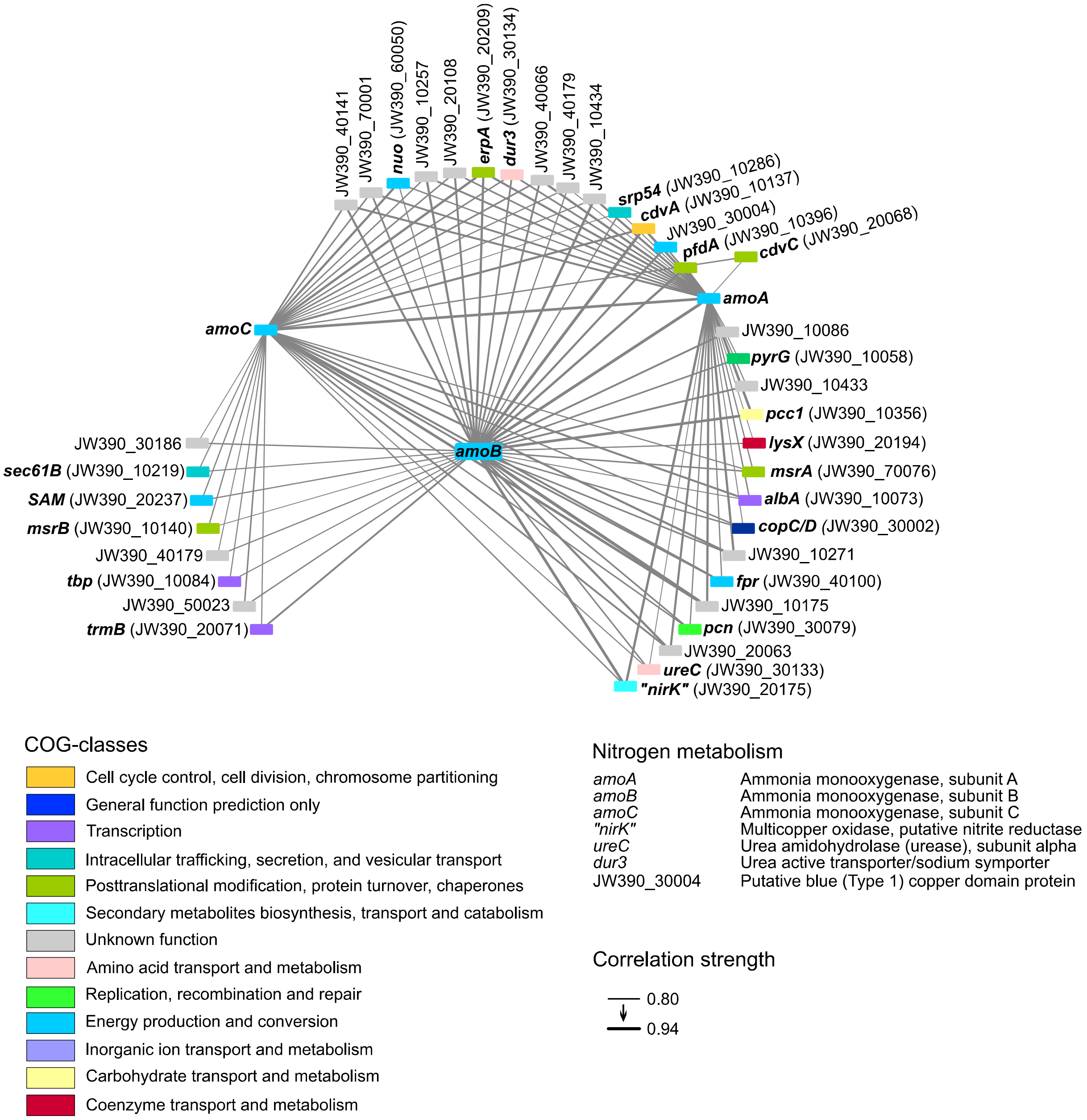


264

265 **Supplementary Figure 12.** Network analysis of ammonia oxidation-related and co-transcribed AOA-
266 LC4 genes over the yearly cycle. Only genes with a strong and significant (Spearman´s $r \geq 0.8$, FDR-
267 corrected p-value <0.05) correlation to at least two of the *amoABC* genes and an average expression
268 higher than the median of all transcribed genes were considered for the network analysis.
269 Correlation coefficient´s strength is visualized by increasing width of the edges. Detailed gene
270 annotation and transcription values can be found in Supplementary Table 2.

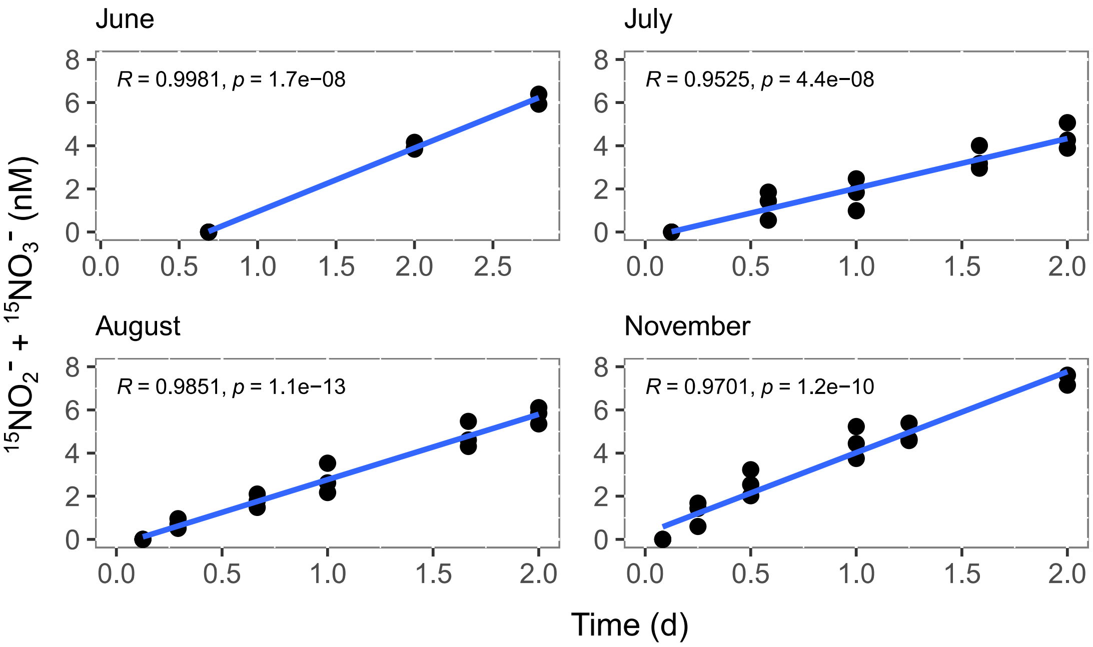


**Supplementary Figure 13.** $^{15}$N-nitrite/nitrate production from $^{15}$N-ammonium in incubations of hypolimnetic water taken from 85 m depth. Produced $^{15}$N-nitrite/nitrate is shown in relation to the first sampling time point, which was set to zero. Linear regression through the time points was used to infer ammonia oxidation rates. All incubations were performed in biological triplicates at 4°C in the dark over a period of 48 h, except for June with 67 h. Samples were taken in 2019 on June 18th, July 29th, August 28th and November 5th. Incubations started within 1–7 h after sampling.

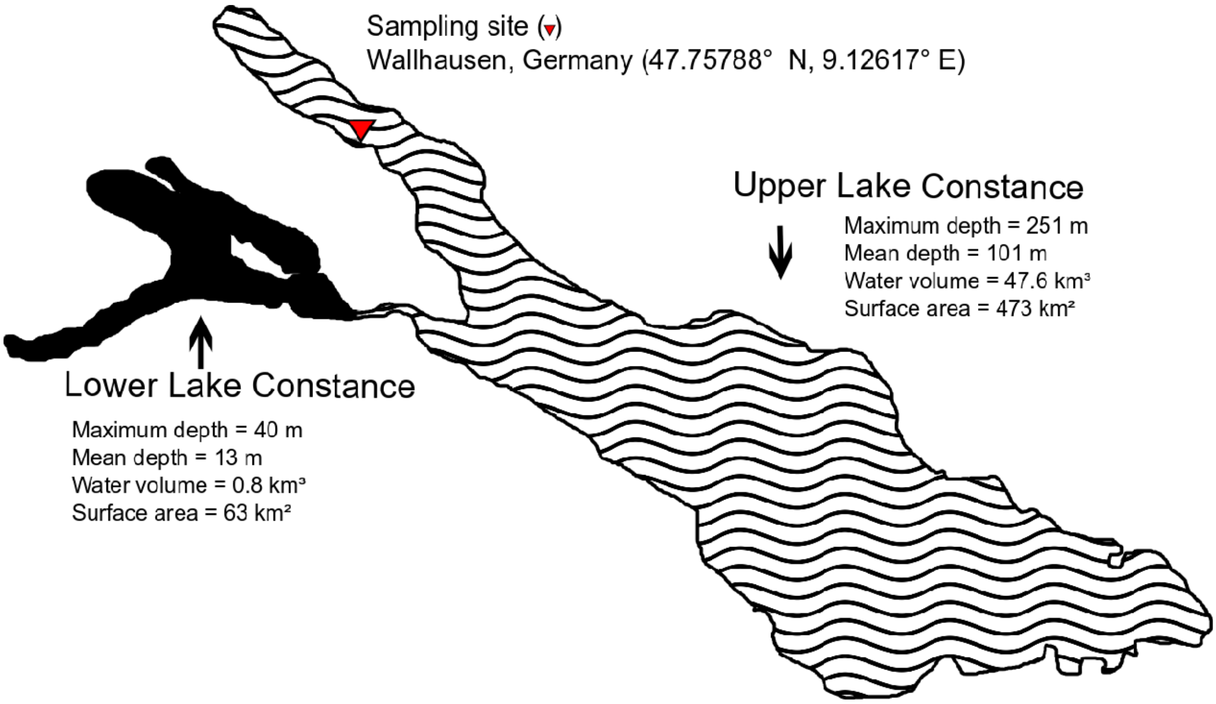


**Supplementary Figure 14.** Map of Lake Constance with the shaded area highlighting Upper Lake Constance as the focus of this study. The sampling site is indicated by a red triangle. All other parameters were taken from Güde and Straile[19].
